# Differential contributions of striatal dopaminergic pathways to visuomotor conditional learning

**DOI:** 10.1101/2024.08.16.608313

**Authors:** Oren Princz-Lebel, Meira M.F. Machado, Miguel Skirzewski, Harleen Rai, Samina Panjwani, Anthony Chu, Claire A. Lemieux, Daniel Palmer, Neeki Derhami, Vania F. Prado, Marco A.M. Prado, Lisa M. Saksida, Timothy J. Bussey

## Abstract

**Rationale:** Midbrain dopamine (DA) neurons project principally towards the striatum, serving as key regulators of movement, motivation, and cognition. Different striatal dopaminergic (DA) pathways may regulate different aspects of cognition.

**Objectives:** We investigated how different striatal DA pathways contribute to a known function of the striatum, visuomotor conditional learning.

**Methods:** Using fiber photometry, we recorded DA transients in these regions as mice learned the touchscreen Visuo-Motor Conditional Learning task.

**Results:** DA transients in all regions dynamically tracked task events, but differed in the timing of peak responses and ramp-like activity preceding a choice, indicating region-specific temporal dynamics across learning. Manipulations of reward probability revealed DA transients in all regions during reward delivery and omission that are consistent with an interpretation in terms of reward prediction error. Thus, DA dynamics in all regions could indicate involvement in visuomotor conditional learning. Therefore, to determine whether nigrostriatal or mesolimbic DA is necessary for learning, we chemogenetically inhibited DA striatal afferents, revealing that only DLS-projecting nigrostriatal DA, and not NAc-projecting mesolimbic striatal DA, was necessary for learning the task.

**Conclusion:** These findings demonstrate functional heterogeneity of aspects of striatal DA signaling, and selective causal roles in the learning of visuomotor conditional learning.

## Introduction

Midbrain dopamine (DA) neurons project principally towards the striatum, serving as key regulators of movement, motivation, and cognition. Given this broad spectrum of influence, it is unsurprising that alterations in striatal DA release play major roles in the neuropathology of several neurodegenerative and neuropsychiatric diseases. For example, the loss of DAergic neurons in the substantia nigra pars compacta is central to Parkinson’s disease pathogenesis (Hirsch et al. 1988; Dauer and Przedborski 2003; Dextera and Jenner 2013; Kalia and Lang 2015), and alterations to DA transmission and receptor expression correspond with the stages and symptom developments of Huntington’s disease (Bernheimer et al. 1973; Spokes 1980; Garrett and Soares-da-Silva 1992).

In rodents, the striatum comprises several interconnected subregions which are thought to be integrated into “limbic” (nucleus accumbens/ ventral striatum, NAc), “associative” (dorsomedial striatum, DMS) and “sensorimotor” (dorsolateral striatum, DLS) circuits (Groenewegen and Uylings 2010). Although these subregions share a common microcircuit architecture with no sharp anatomical boundaries, subtle gradients in receptor density, interneuron proportions and protein expression are believed to introduce heterogeneity across the striatum in terms of both composition and function (Cox and Witten 2019). For example, properties of the DA system across the different regions of the striatum have recently been shown to differ in gene expression (Poulin et al. 2018; Heymann et al. 2020), signaling profiles (Jones et al. 1995; Cragg et al. 2000; Brown et al. 2011; Sulzer et al. 2016), and local mechanisms for modulation of release (Threlfell et al. 2010, 2012a).

This anatomical and neurochemical heterogeneity is increasingly believed to underlie functional specialization across striatal dopaminergic circuits. Consistent with this view, regional differences in dopamine function have been reported across multiple cognitive domains. For example, amplifying DA signals in the DMS accelerates the emergence of compulsive behaviour, while the same manipulation in DLS does not (Seiler et al. 2022a). Similarly, DA-deficient mice with DA levels restored only in DLS showed intact object recognition, spatial memory, discriminatory and visuospatial learning, yet remain impaired in reward-mediated lever pressing, indicating that distinct behavioural processes depend on different striatal dopaminergic circuits (Darvas and Palmiter 2009). Furthermore, dopamine signals in the NAc update more rapidly than those observed in the DLS during stimulus-reward learning, highlighting region-specific contributions to adaptive decision-making (Versprille et al. 2026). Collectively, these studies suggest that dopamine may serve different functions, depending on region.

One striatal-dependent cognitive domain for which the contribution of dopamine circuit heterogeneity remains poorly understood is stimulus-response (S-R) learning. This form of associative learning underlies the ability to form and maintain automatic behaviors, linking environmental stimuli with corresponding motor responses. A fundamental cognitive process, S–R learning is critical for adaptive decision-making and efficient execution of routine actions (Jog et al. 1999; Barnes et al. 2005; Yin et al. 2009; Thorn et al. 2010; Liljeholm and O’Doherty 2012; Smith and Graybiel 2013; Cox and Witten 2019b). Notably, aberrant S–R learning is frequently observed in conditions characterized by altered dopaminergic transmission, including Parkinson’s disease (Hadj-Bouziane and Boussaoud 2003; Hayes et al. 2015; Wu et al. 2015; Olson et al. 2019; Mi et al. 2021) and Huntington’s disease (Knopman and Nissen 1991; Lawrence et al. 1999), underscoring the central role of dopamine in modulating the learning and expression of associative behaviors, and the relevance of S-R learning to understanding neurodegenerative and neuropsychiatric disease. However, evidence linking dopamine to S-R learning has largely arisen from disease-associated dysfunction, lesion studies, or broad manipulations of dopaminergic transmission. As a result, the causal contribution of anatomically distinct striatal dopaminergic circuits to S-R learning remains unresolved. In particular, it is unknown whether the functional specialization observed across striatal dopamine systems in reward learning, decision-making, and compulsive behaviour also applies to the acquisition and performance of S-R associations. Addressing this question is critical for understanding both the neural basis of associative learning and how dopaminergic dysfunction contributes to cognitive deficits in neurological disease.

In the present study, therefore, we first capitalized on recent technological advances in optical imaging methods combined with genetically encoded biosensors (Sun et al. 2020; Wu et al. 2022; Piantadosi et al. 2025) to evaluate the possible contributions of dopamine in different compartments of the striatum –NAc, DMS and DLS --during the learning of the of striatal-dependent ‘Visuomotor Conditional Learning’ (VMCL) task (Horner et al. 2013; Delotterie et al. 2014), designed to assess S-R learning. We found differences in dopamine dynamics across regions, but also signals consistent with a role in learning in all regions. Therefore, to determine whether nigrostriatal or mesolimbic DA was necessary for learning, we chemogenetically inhibited DA striatal afferents, revealing that only DLS-projecting nigrostriatal DA, and not NAc-projecting mesolimbic DA, was necessary for learning the task. Our findings clearly show a role for the DLS-projecting nigrostriatal, but not the NAc-projecting mesolimbic, dopamine pathway in visuomotor conditional learning.

## Materials and Methods

### Animals

Male and female adult (8-week-old) wild-type C57BL/6J mice were obtained from Jackson Laboratories (strain #000664, Bar Harbor, ME). Heterozygous male and female DAT-IRES-Cre mice (Slc6a3Cre) were backcrossed with C57BL/6J mice for at least 8 generations and maintained as an inbred strain in our mouse colony. All procedures were conducted in accordance with the guidelines of the Canadian Council of Animal Care guidelines and approved by the Animal Care and Veterinary Services (ACVS) from Western University (protocols # 2017-031, 2021-082, 2020-163) and in compliance with the ARRIVE guidelines.

### Experimental Design

Experiments began when mice were 2- to 4-months of age. Animals were housed in 28 x 18cm plastic shoebox cages with wire tops at 22-23°C, 50 ±10 % humidity, with a 12:12h reverse light-dark cycle (lights turned off at 09:00). Each cage was supplied with Biofresh bedding, Envirodri, and Nestlets nesting materials, along with twist bits, diamond twists, wooden chew sticks and a cardboard tunnel for environmental enrichment. Food was provided *ad libitum* until 1-2 week(s) prior to behavioral testing, at which point mice were food restricted to 85-90% of their free-feeding body weight. Experiments were performed during the dark cycle (between 09:00 and 17:00) 4-7 days/ week and mice were weighed and fed directly upon return to the home cage after each session. Prior to stereotaxic surgery, mice were housed in groups of 2 to 4. Following surgical procedures, mice were single housed for the remainder of the experiment to prevent infection of the surrounding incision area or damage to the fiber optic implant.

To characterize how DA neuromodulation across the striatum impacts visuomotor conditional learning, independent cohorts of wild-type or mutant mice performed the automated touchscreen VMCL task. Separate cohorts of mice were used for unilateral fiber photometry recordings in the NAc (*n*=12), DMS (*n* =12), and DLS (*n* =9), and bilateral chemogenetic inhibition for nigrostriatal DA projections (*n* =15), mesolimbic DA projections (*n* =17) and their corresponding controls (nigrostriatal control *n* =18; mesolimbic *n* =19).

### Automated Touchscreen Cognitive Assessment

#### Apparatus

Cognitive testing was conducted using automated Bussey-Saksida Mouse Touchscreen Systems Model 80614-20 (Lafayette Instruments, Lafayette, IN). Mice were assessed on the touchscreen VMCL task as previously described (Delotterie et al. 2014). Experiments were conducted inside sound-attenuating cabinets containing an operant chamber and a touch-sensitive 12.1-inch monitor (screen resolution 600 x 800). The operant chamber was trapezoidal-shaped and constructed from three black Plexiglas walls, which open to the touchscreen (20 cm high X 18 cm long X 24 cm) (Fig. 1A). The ceiling of the chamber was made of clear Plexiglas, and the floor consisted of perforated stainless steel with a waste tray situated below. Throughout training, the touchscreen was permanently covered by a black Plexiglas 3-holed mask, which limited subjects’ interaction with the screen to relevant task locations (three 7×7cm square windows). The chamber was equipped with a liquid reward dispensing magazine situated at the back of the chamber, linked to a liquid reward dispenser pump (reinforcer: strawberry milkshake, Neilson Dairy). A light-emitting diode (LED) illuminated the food magazine during reward delivery and a tone generator was installed that triggered auditory tones used throughout touchscreen training. Animal activity was recorded via infrared photobeams located at the front and back of the chamber and at the entry to the reward magazine. The schedule design, control of the apparatus via Whisker control system and data collection was controlled by ABET II Video Touch software V21.02.26.

**Fig. 1.**
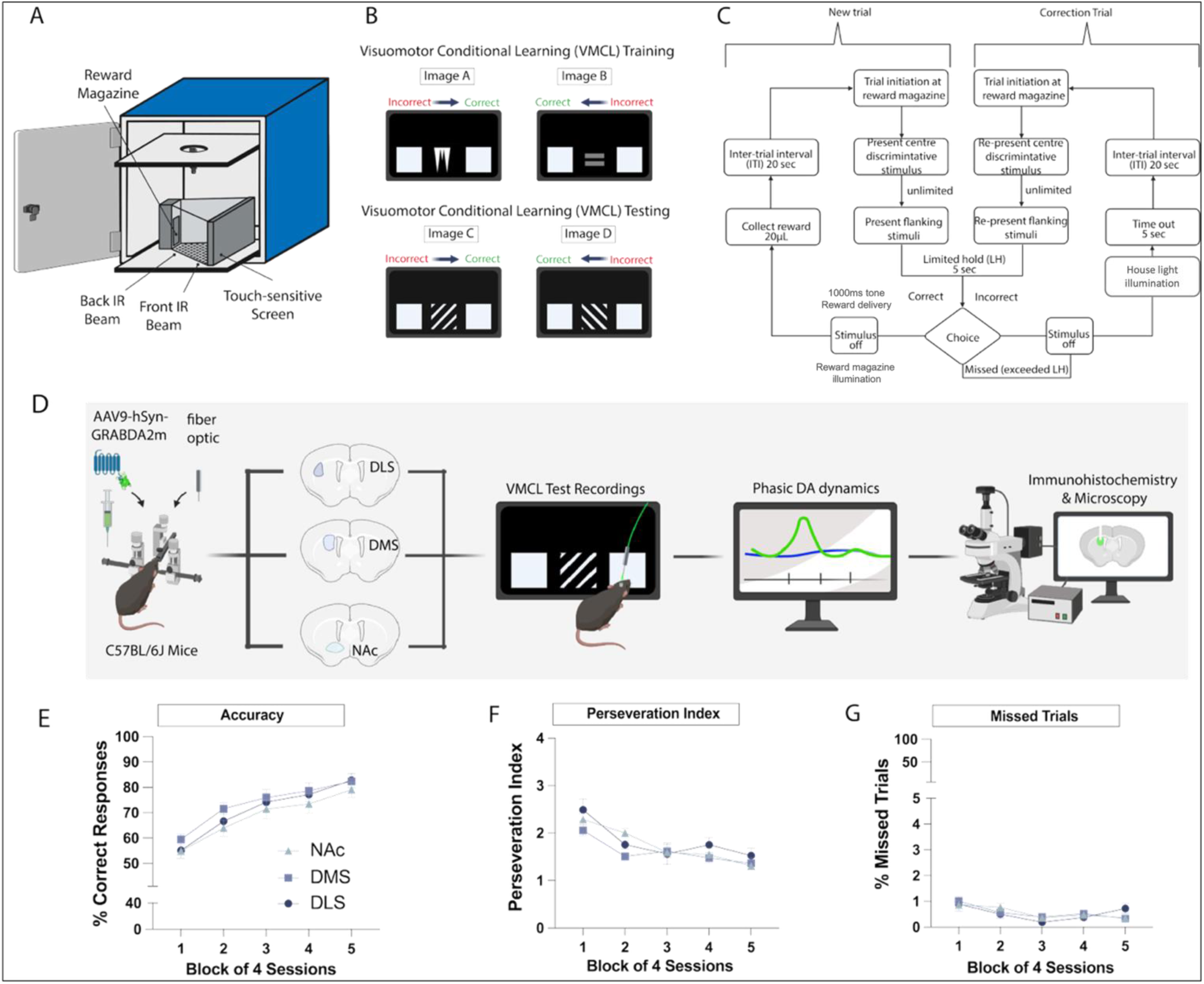
The Visuomotor Conditional Learning (VMCL) task for assessing visuomotor conditional learning. (A) Behavioural assessments were conducted within touchscreen systems equipped with a touch-sensitive screen, a reward magazine attached to a reward pump for delivery of strawberry milkshake liquid reward, infra-red beams in the reward magazine and front and back chamber and ABET cognition software. (B) The VMCL task was designed to evaluate S-R learning and expression, with subjects learning the conditional rule: “if visual stimulus A is presented, make motor response X; if visual stimulus B is presented, make motor response Y”. All subjects underwent VMCL training prior to VMCL testing for 20 sessions, 5-6 sessions/week. (C) Flowchart overview of the VMCL task. Following trial initiation, one of two discriminatory stimuli was presented. Touching it resulted in the presentation of two choice stimuli in the flanking locations (left and right), for a 5-second limited hold period. A correct choice triggered the illumination of the reward magazine light, the delivery of 20uL of strawberry milkshake, and a 1000ms tone. The reward collection was followed by an intertrial interval (ITI) of 20 seconds. An incorrect choice or missed trial was followed by a time-out period, indicated by illumination of the house light for 5 seconds, and an ITI of 20 seconds with the repetition of this same trial (i.e. a correction trial (CT)). The CT loop continued until a correct response was made. (D) Experimental design of recording *in vivo* DA dynamics in freely moving animals as they performed the VMCL task. Wildtype C57BL/6J mice underwent stereotaxic surgery for viral injection of AAV9-hSyn-GRABDA2M into either the nucleus accumbens (n=12), dorsomedial striatum (n=12) or dorsolateral striatum (n=9), and surgical implant of a fiber optic probe. Following post-operative recovery, mice ran on the VMCL Test for 20 sessions during simultaneously recording with fiber photometry. No significant differences were found between groups in VMCL test (E) accuracy, (F) perseveration index (G) or number of missed trials. Data presented as Mean + SEM, group x session Two-way RM ANOVA or RM Mixed-Effect Models, p>0.05.

#### Touchscreen habituation and pre-training procedure

Mice were habituated to the strawberry milkshake reinforcer and the touchscreen apparatus at the beginning of the touchscreen-based behavioral testing. The stages of acclimation and appetitive conditioning were conducted as previously described (Mar et al. 2013; Delotterie et al. 2014) and can be found in full within the standard operating procedure freely available at touchscreencognition.org. In brief, food-restricted mice were first acclimated to the strawberry milkshake reward in their home cage for 2-3 consecutive days. Next, mice were assigned to a touchscreen operant chamber and gradually exposed to the apparatus for 10-40 minutes per session, along with habituation to the strawberry milkshake reward delivery and associated tone cue (1 second, 3kHz, 80 dB). A maximum of one session occurred each day.

Once acclimated to collecting reward within the reward receptacle, mice progressed to several sequential pre-training procedures wherein they learnt to (a) detect and respond (usually through nose pokes) to a training stimulus that appeared in one of the 3 touchscreen windows for 20µL of strawberry milkshake reward, and (b) initiate a trial by nose poking the reward receptacle following an intertrial interval (ITI) (20 seconds ITI). These pretraining sessions were repeated until each mouse reached a criterion of 30 completed trials within a 60-minute session. Each new trial and correction trial began in the reward receptacle, as the mouse enters the reward magazine and thus breaks the photobeam across the entry point of the magazine.

Finally, the last two training schedules prepared the mice for key procedures that are fundamental to the VMCL task. For example, mice first learnt to selectively respond to a target training image, with a 5 second time-out punishment if an incorrect touch occurred. Once this was accomplished, mice also learnt to nose poke the screen twice: first touching a training stimulus in the central location, and then touching a training stimulus in the left or right flanking windows to receive a reward within a decreasing time window (this limited hold period began at 10 seconds and then was reduced to 5 seconds). Both procedures were included in the design of the VMCL task to promote immediate and rapid S-R-driven behavior. These stages were repeated consecutively until mice reached a criterion of 23/30 correct trials within a 60-minute session for two consecutive sessions. Upon successful pretraining completion, subjects were given free food and prepped for stereotaxic surgery.

#### Visuomotor Conditional Learning (VMCL) training task (Image set A+B)

Following stereotaxic surgery and recovery (see below), subjects were re-established on food restriction, re-baselined on criterion for the last pretraining stage and then advanced to the VMCL task training.

Each session began with a 20µL strawberry milkshake reward (800ms pulse feed time) delivered to the reward receptacle at the back of the chamber. When the mouse exited the receptacle, a trial began with the presentation of one of two discriminatory visual stimuli in the centre window: a white icicle (image A) or a grey equal sign (image B). Through trial and error, subjects learnt that the presentation of the white icicle (image A) required an immediate nose poke to the right flanking window for a correct response and the delivery of reward. In contrast, the presentation of the grey equal sign (image B) required a nose poke to the left flanking window for a correct response and delivery of the reward. Both discriminatory images were presented equally throughout the session (15 trials of each, for a total of 30 new trials maximum), and were presented pseudo-randomly so that no image was presented more than 3 times in a row.

Although the time to nose poke the discriminatory image in the central window was uncapped, the time to respond to the left or right flanking window was limited to 5 seconds to promote fast and automatic S-R-driven behavior (5-second limited hold period). A correct response immediately triggered the illumination of the reward tray light, the presentation of 20uL of strawberry milkshake, and a 1000ms tone, followed by an intertrial interval (ITI) of 20 seconds once the reward was collected. An incorrect response, or a failure to respond within the 5-second limited hold period (termed a ‘missed’ trial) led to a time-out period, indicated by illumination of the house light for 5 seconds, and followed by an ITI. To counteract the development of side biases and ensure that subjects received a consistent number of rewards per session, any trial immediately succeeding a time-out period was designated a ‘correction trial’ in which the same image was re-presented. There was no limit to the number of correction trials that could be presented consecutively, but once a correct choice was made, the correction procedure ended, and normal non-correction trials resumed. VMCL training involved a maximum of 30 new (non-correction) trials in each maximum 60-minute session. Each new trial and correction trial began in the reward receptacle, as the mouse enters the reward magazine and thus breaks the photobeam across the entry point of the magazine.

For fiber photometry experiments, VMCL training occurred predominantly with mice untethered and without *in vivo* recordings. Well-trained mice were subsequently acclimated to the fiber optic tether in the last few sessions of VMCL training. For chemogenetic inhibition experiments, the final sessions of VMCL training were used to test whether acute doses of the chemogenetic agonist clozapine N-oxide (CNO) or saline control (1 mg/kg, intraperitoneally (*i.p.)*; order counterbalanced) would impair well-trained mice (>77% accuracy over two consecutive days) on the expression of previous learning.

#### Visuomotor Conditional Learning (VMCL) testing task (image set C+D)

VMCL testing procedures were identical to VMCL training, except new discriminatory stimuli were used: right-leaning and left-leaning diagonal lines of the same luminance and size. The visual stimulus with the right-leaning diagonal lines (image C) was rewarded with 20uL of strawberry milkshake only when the right flanking window was nose-poked, whereas the visual stimulus with the left-leaning diagonal lines (image D) was rewarded only when the left flanking window was nose-poked. Similar to VMCL training, VMCL testing involved a maximum of 30 new (non-correction) trials in each 60-minute capped session. For fiber photometry experiments, mice were tethered and recorded every day during VMCL testing for 20 consecutive sessions to ascertain how striatal DA transients respond and evolve throughout the learning and the expression of learning. For chemogenetic inhibition experiments, mice received systemic injections of CNO (1mg/kg, *i.p.*) 5 days a week for 20 sessions to assess how nigrostriatal or mesolimbic DA blockade impacts new learning. Additionally, following learning, mice received 4 more sessions of saline injections, and then 4 additional sessions of CNO injections (same dose) to explore the impact of inhibition relief and reinstatement, respectively.

#### Visuomotor Conditional Learning (VMCL) Testing Probes

In a subset of animals undergoing striatal DA fiber photometry recordings, learning of VMCL was followed by a set of behavioral probes to explore how subjects and their associated striatal DA transients responded to changes in reward probability following learning. All subjects were well-trained on VMCL prior to probe testing, and each behavioral probe was preceded by a re-baselining session to ensure high accuracy. Furthermore, correction trial loops were disabled for all behavioral probes.

##### Visuomotor Conditional Learning (VMCL) Testing 80-20 Probability Probe

The first VMCL Testing Probe conducted was an 80-20 probability probe. In comparison to VMCL Testing during which correct and incorrect choices were followed by 100% and 0% reward probability, respectively, during the probability probe, correct and incorrect choices were followed by 80% and 20% reward probability, respectively. This provided us with 4 balanced trial conditions at choice: (a) *correct rewarded*, (b) *correct not rewarded*, (c) *incorrect rewarded* and (d) *incorrect not rewarded*. Subjects had a maximum of 30 trials in each 60-minute capped session, for 5 consecutive sessions.

##### Visuomotor Conditional Learning (VMCL) Testing 70-30 Probability Probe

Following completion of the 80-20 probability probe, subjects were re-baselined and then tested on a 70-30 probability probe. Here, the setup mirrored the 80-20 probability probe except the probability of receiving a reward following correct choices was set at 70% and following incorrect choices at 30%. Subjects had a maximum of 30 trials in each 60-minute capped session, for 5 consecutive sessions.

### Behavioral Data analysis

All behavioral analyses were imported to and conducted on GraphPad Prism 9 for macOS Version 9.5.1 (528). The primary outcomes of interest for VMCL behavior were (a) **accuracy** (percentage of correct responses) which was calculated by the equation 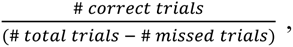, (b) **percentage of missed trials**, which was calculated by the equation 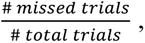, (c) **number of correction trials**, (d) **perseveration index** (a measure of repeated incorrect responding during correction trial loops that corrects for the number of opportunities to do so), which was calculated by the equation 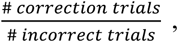, (e) **correct touch latency** (time from nose poking the discriminatory stimulus in the central window to nose poking the correct right or left flanking window), (f) **incorrect touch latency** (time from nose poking the discriminatory stimulus in the central window to nose poking the incorrect right or left flanking window), and (g) **reward collection latency** (time to nose poke into the reward magazine after giving a correct response on the right or left flanking window of the touchscreen). Unless otherwise denoted, all behavioral data were expressed as means + SEM, and plotted in 5 blocks of 4 sessions. Shapiro-Wilk test of normality and the Rout method for identifying outliers (Q=0.1%) were conducted prior to each analysis. If normality failed, data were transformed prior to statistical testing[40]. Differences between two means were examined using paired or unpaired t-tests. When examining 3 or more comparisons, tests of analysis of variance (ANOVA) were employed (one- or two-way with repeated measures), followed by Šídák’s multiple comparisons if a significant interaction was revealed. Alternatively, if a dataset had missing values due to the identification of outliers, we compared groups using a repeated measures mixed-effects ANOVA model. Estimation of sample size in VMCL task experiments was estimated using a partial eta squared (ηp 2) power analysis for repeated measures two-way ANOVA (power = 0.8, ɑ = 0.05)[41]. For experiments combining fiber photometry and touchscreens, we used standard sample sizes (N = 9–12) as previously reported (Markowitz et al. 2018; Legaria et al. 2022; Skirzewski et al. 2022). In all experiments, both male and female mice were tested in balanced design (Galea et al. 2020), but were combined in statistical analyses due to insufficient power to detect sex differences (Miller et al. 2017). Analyses were considered significant if p values (or adjusted p values) were < 0.05.

### Viral vectors

For fiber photometry experiments, recording extracellular DA across the striatum was achieved by injecting 500nl of AAV9-hSyn-GRABDA2m (3.1 × 1013gc/mL, Vigene Biosciences) unilaterally into the NAc, DMS or DLS. For chemogenetic inhibition of nigrostriatal or mesolimbic DA afferents, expression of hM4D(Gi) was achieved by injecting 400nl of the retrograde AAVrg-hSyn-DIO-hM4D(Gi)-mCherry (8 x 1012 gc/mL, Addgene, #44362-AAVrg) bilaterally into the DLS or NAc of DAT-IRES-Cre heterozygous mice, respectively. To control for the known side effects of CNO[47,48], we created a littermate control group lacking hM4D(Gi), by injecting the retrograde AAVrg-hSyn-DIO-mCherry (7 x 1012 gc/mL, Addgene, #50459-AAVrg) into the DLS or NAc in DAT-IRES-Cre heterozygous mice.

### Histology

Mice were anesthetized with ketamine (100 mg.kg-1)-xylazine (20mg.kg-1) and transcardially perfused with ice-cold phosphate-buffered saline (PBS) followed by 4% paraformaldehyde (PFA). Brains were extracted and kept overnight in 4% PFA at 4°C, and then transferred into 30% sucrose solution where they stayed until they were flash-frozen on dry-ice in Tissue-Tek O.C.T. Compound (Sakura, Cat. #4583) for long-term storage at -80°C. Tissue was sliced on a cryostat (Leica CM1950s) or vibratome (Campden Instruments 5100mz) in 50µm coronal sections. Dorsal and ventral striatum (AP 1.9 to -1.9) and/or VTA/SNc (AP -2.4 to -4.0) were identified using a mouse brain atlas (Paxinos & Franklin, Academic Press, 2001) and appropriate slices collected in a PBS with 0.02% sodium azide solution.

### Immunohistochemistry

#### GFP stain for confirmation of GRABDA2M expression

Sections were mounted onto SuperFrost+ slides and surrounded by a hydrophobic barrier using a Pap Pen (ThermoFisher, Cat. #R3777). In a humid chamber, sections were rinsed twice with Tris-buffered saline (TBS) for 5 minutes and then incubated in TBS containing 1.2% Triton X-100 for 20 minutes. The sections were rinsed with TBS for 10 minutes and then blocked in TBS containing 5% (v/v) normal goat serum at room temperature for 1 hour. After blocking, sections were rinsed twice in TBS for 10 minutes and then incubated overnight at 4 °C with chicken anti-GFP (Abcam, ab13970, 1:500) in TBS containing 0.2% Triton X-100 and 2% normal goat serum. The following day, sections were washed twice in TBS for 10 minutes and incubated for 1 hour with Alexa 488 goat anti-chicken (Thermo Fisher, A11039, 1:500) in TBS 0.2% Triton X-100 and 2% normal goat serum. The sections were washed twice in TBS for 10 minutes and then incubated with Hoechst 33342 (Thermo Fisher H3570, 1:1000) for 5 minutes to counterstain nuclei. Images were captured using the Leica DM6B Thunder imager (Leica Microsystems Inc.).

#### mCherry stain for confirmation of chemogenetic expression + colocalization with DA neurons expressing tyrosine hydroxylase (TH)

Protocol mirrored the one above, except the primary and secondary antibodies differed: primaries were chicken anti-TH (Abcam, ab76442, 1:500) and rabbit anti-mCherry (Abcam, ab167453, 1:1000) and secondaries were Alexa 488 goat anti-chicken (Invitrogen, A11039, 1:500) and Alex 594 goat anti-rabbit (Invitrogen, A11012, 1:500). Images were captured using the Stellaris 5 Confocal Microscope (Leica Microsystems Inc.).

#### c-Fos stain for confirmation of chemogenetic inhibition

Protocol is similar to the one above, but before the first washes, sections were incubated with citrate buffer (sodium citrate 10mM, 0.05% Tween, pH 6.0) at 95°C for 20 minutes for antigen retrieval and cooled down at room temperature for 30 minutes on ice. The primary antibodies were guinea pig anti-c-Fos (Synaptic Systems, 226-308, 1:1000), rat anti-mCherry (Invitrogen, M11217, 1:1000) and chicken anti-tyrosine hydroxylase (TH, Abcam, ab76442, 1:1000). Secondary antibodies were Alexa 488 goat anti-guinea pig (Invitrogen, A11073, 1:500), Alexa 594 anti-rat (Invitrogen, A11007, 1:500), and Alexa 647 anti-chicken (Invitrogen, A21449, 1:500). Images were captured using the Leica Stellaris STEM Confocal Microscope (Leica Microsystems Inc.). Cells were quantified using ImageJ software (National Institutes of Health). Data were shown as the percentage of mCherry colocalized cells for VTA and SN. In the DSL and NAc, data were shown as the percentage of total c-Fos positive cells.

### Surgical procedures for fiber photometry recordings

Stereotaxic surgery for viral infusions and fiber optic implantations were executed as previously described (Kljakic et al. 2022a; Skirzewski et al. 2022). In brief, mice received an injectable dose of meloxicam (5mg/mL, *i.p.)*15 minutes prior to surgery. Mice were anesthetized with a 4% isoflurane induction rate and placed in a stereotaxic frame, where they were maintained at 1.5-3%. A heating pad and rectal thermometer were used to maintain body temperature throughout the entirety of the surgery. Following fur removal and aseptic procedures, the top of the skull was exposed, and holes were drilled for the viral infusion needle, fiber optic implant and two skull screws. Viral injections into the striatum were made with a micro syringe pump (0.5µL, 0.25µL/min) at coordinates (**NAc**= AP: 1.8mm, ML: 0.5mm, DV: 4.0mm; **DMS**= AP: 0.75, ML: 1.3 ML, DV: 2.5; **DLS**= AP: 0.75, ML: 2.3, DV: 2.5) from bregma. Injectors were left in place for 5 minutes after infusion and slowly removed. For all fiber photometry recordings, mice were injected unilaterally, the hemisphere counterbalanced across subjects within each experimental group. Low-auto-florescence fiber optic implants (400µm O.D. 0.48 NA, 4.5mm-long, made in-house) were inserted 0.1mm above injection site and chronically implanted with dental cement (Stoelting 51458). After surgery, mice were allowed to recover on a heating pad until ambulatory, then returned to their homecage with moistened chow available. Mice then underwent a 3-week recovery period to allow for viral expression.

Mice were acclimated to the fiber patch-cord tethering during VMCL training (see above). To record fluorescence signals, a Doric Lenses photometry system was used, equipped with a fluorescent mini-cube which transmitted sinusoidal 465 nm LED light modulated at 572 Hz and a 405 nm LED light modulated at 209 Hz. LED power was set at approximately 25µW. Florescence was collected through a patch-cord connected to the fiber optic implant of each mouse and transmitted back to the mini-cube, amplified, and focused into an integrated high sensitivity photoreceiver (Doric Lenses). Real-time florescent signal was sampled at 12 kHz and demodulated and decimated to 100 Hz using Doric Studio Software V5.2.2.3. To time-lock florescence signals with the occurrence of behavioral events in the touchscreen, a transistor-to-transistor logic (TTL) pulse was sent from ABET II (Lafayette Instruments) at task initiation. All behavioral measures (i.e. trial initiation, correct choice, incorrect choice, or reward collection) were generated by the ABET II software.

### Fiber photometry analysis and statistics

Analyses of fiber photometry signals were done with a custom-written publicly available python script (https://doi.org/10.5281/zenodo.7577053), as previously described (Piantadosi et al. 2025). Fluorescent signals from 405 and 465 nm channels were low band-pass filtered to remove events exceeding 3 Hz. The isosbestic 405 nm channel was exploited to correct for bleaching or movement artifacts. A least-squares linear fit model was applied to isosbestic 405 nm signal to align it to the 465 nm signal, producing a fitted 405 nm signal used to normalize the 465 nm as follows: 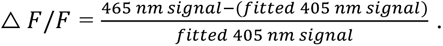. This signal was then normalized into a z-score, where z-score was calculated as follows: z-score= △ *F*/*F* –(µbaseline)/ (σbaseline). µbaseline is the mean △ *F*/*F* from baseline period, which was taken from the continuous signals in the last 10 seconds of the ITI, based on assessments of signal variance across non-event-related time periods. σbaseline is the standard deviation *F*/*F* from the same baseline period. GRABDA2M signals were filtered and aligned to various task events (e.g. trial initiation, correct choice, incorrect choice and reward collection) to assess how changes in fluorescence (ΔF/F) track task performance.

Validation of the *in vivo* DA dynamics using GRABDA2M and fiber photometry was achieved by recording in a touchscreen apparatus whose screen was turned off, with △ *F*/*F* fluorescence baseline signaling during 5 min before administering systemic saline injections (*i.p.)* for a 1h long recording. Finally, mice received an injection of cocaine (10mg/kg, *i.p.*) and florescence was recorded for another hour.

Line graphs illustrating z-scores from GRABDA2M signalling consisted of averaged trials per session for each mouse and then averaged all subjects within the same group (NAc-probe implanted, DMS-probe implanted, or DLS-probe implanted) and were generated on GraphPad Prism V10.1.2 for Mac (GraphPad Software, San Diego, CA). Each began with 10 seconds prior to the task event of interest and ended 10 seconds following the event of interest. The generation of heatmaps, estimation of area under the curve (AUC), and max peak analysis of events were obtained using OriginPro 2021 V9.8.0.200 (OriginLab Corporation, Northampton, MA). Briefly, heatmaps illustrating z-scores from GRABDA2M signalling were constructed of individual trials of represented animals. Each began with 10 seconds prior to the task event of interest and ended 10 seconds following the event of interest. To find peaks, calculate height, and to integrate the AUC, the averaged trial signal from each session across animals was used to find and calculate peaks (curve) within the region of interest. This allowed us to obtain a baseline from Y=0 within the window search, which corresponded with a z-score deviation of 0. AUC and max peak response following task events (e.g. trial initiation, correct and incorrect choice and reward collection) were then calculated by setting the window search to the 10s following task onset. Alternatively, baseline or ramp analyses were calculated by setting the window search from -10 seconds to -5 seconds prior to event onset, or -5 seconds to 0 seconds prior to task onset, respectively. All data were imported to GraphPad Prism V10.1.2 for Mac (GraphPad Software, San Diego, CA) for statistical analysis.

### Surgical procedures for chemogenetic inhibition

Mice received an injectable dose of meloxicam (5mg/mL) *i.p.*, 15 minutes prior to surgery. Mice were anesthetised with a 4% isoflurane induction rate and placed in a stereotaxic frame, where they were maintained at 1.5-3%. A heating pad and rectal thermometer were used to maintain body temperature throughout the entirety of the surgery. Following fur removal and aseptic procedures, the top of the skull was exposed, and holes were drilled to accommodate insertion of the viral infusion needle. Viral injections (see viral vectors section above) into the striatum were made with a microsyringe pump (0.5µL, 0.1µL/min) at coordinates (**NAc**= AP: 1.8mm, ML: 0.5mm, DV: 4.0mm OR **DLS**= AP: 0.75, ML: 2.3, DV: 2.5) from bregma. Injectors were left in place for 5 minutes after infusion and slowly removed. For all chemogenetic inhibition experiments, mice were injected bilaterally. Following viral infusions, incisions were closed using simple-interrupted suturing with absorbable sutures (Ethicon, Monocryl 5-0, Y433H). After surgery, mice were allowed to recover on a heating pad until ambulatory, then returned to their homecage with moistened chow available. Mice then underwent a 3-week recovery period to allow for viral expression.

## Results

### *In vivo* dopamine dynamics across the striatum track VMCL task events

The VMCL task (Fig. 1B-C) is a well-established instrumental paradigm designed to assess the learning and expression of S-R contingencies through the repeated pairing of visual stimuli (presented on a touch-sensitive screen) with arbitrary motor responses (nose pokes to the right- or left-flanking positions). The VMCL task was designed to be solvable by establishing distinct S-R associations, rather than action-outcome strategies or by simple Pavlovian conditioning processes alone (Reading et al. 1991), and has previously been shown to be dependent on the striatum, and not other related structures such as the hippocampus (Delotterie et al. 2015).

First, we verified that wild-type C57BL/6J mice that underwent stereotaxic surgery for *in vivo* fiber photometry recordings in the NAc (*N* =12, *n*=6^♂^, *n*=6^♀^), DMS (*N*=12, *n*=5^♂^, *n*=7^♀^), and DLS (*N*=9, *n*=5^♂^, *n*=4^♀^) acquired the task at a similar rate. While tethered, the percent correct accuracy of all 3 groups increased across sessions but there was no main effect of group or interaction (Fig. 1E; 2-way Repeated Measures [RM] ANOVA –main effect of session: F3.125,87.49= 54.60, p<0.0001). Furthermore, all subjects reached asymptote –defined as no difference between consecutive mean percent correct scores, and the point at which behavior might be expected to have become habitual –in the last four sessions of the VMCL task, with no group differences (Fig. 2D). Together, these comparisons indicate that our mice acquired the VMCL task to stable asymptotic levels of performance, irrespective of the site of probe implantation. These findings provide confidence that any observed differences between the NAc, DMS or DLS DA dynamics described below were not simply a result of behavioral differences across groups.

**Fig. 2.**
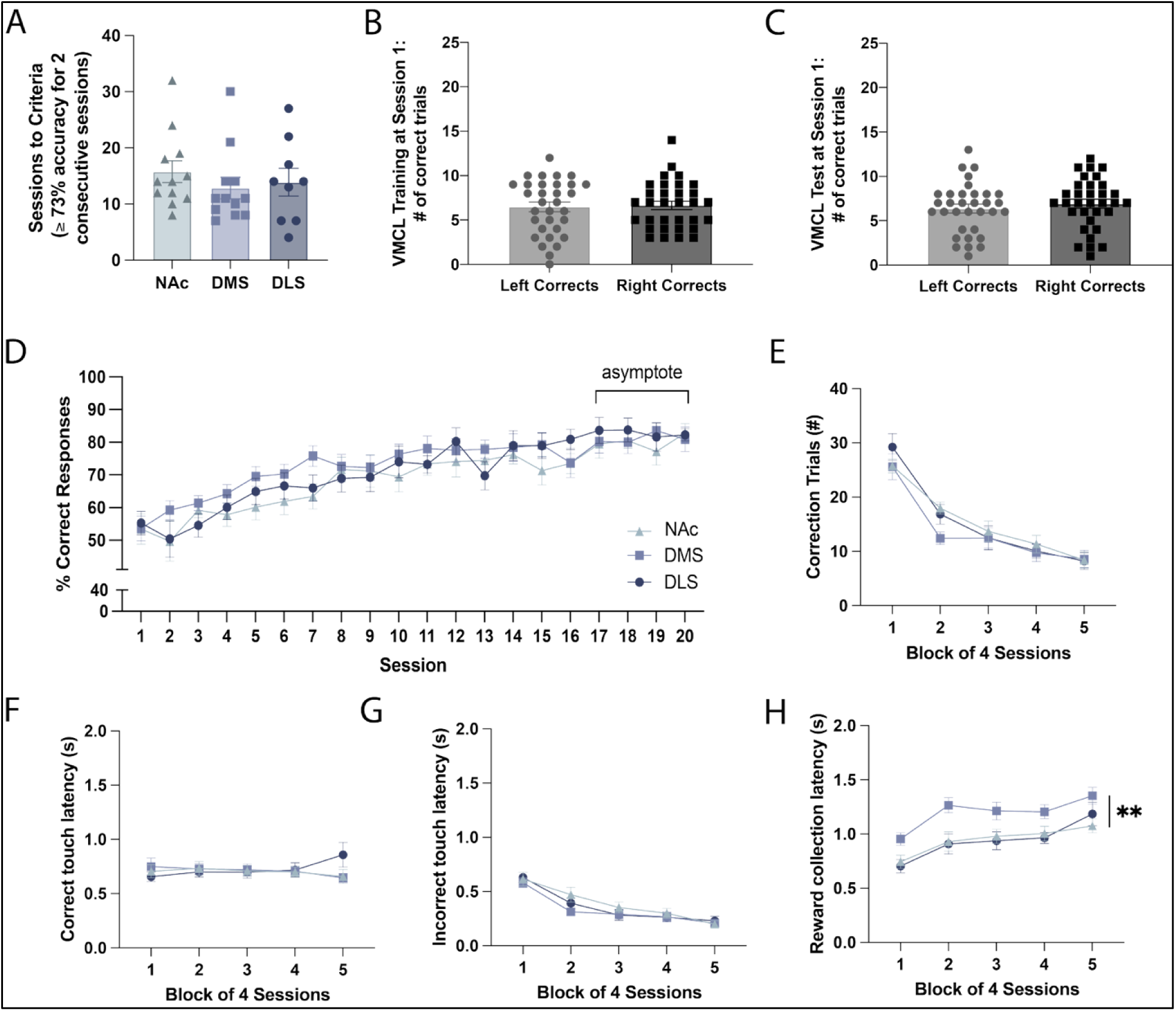
Minimal differences between nucleus accumbens-, dorsomedial- and dorsolateral striatum-probe-implanted mice on secondary measures of behaviour during the visuomotor conditional learning task. C57BL/6J mice that underwent stereotaxic surgery for *in vivo* fiber photometry recordings in the nucleus accumbens (NAc), dorsomedial striatum (DMS) and dorsolateral striatum (DLS) showed no significant differences in the sessions to reach criterion in the visuomotor conditional learning (VMCL) training task (≥73% accuracy for two consecutive sessions) (A). Furthermore, task performance began at chance levels, with no significant differences between accuracy on the two arms of the conditional rule (responses to the left-flanking or right-flanking positions) in the VMCL training task (B) or VMCL test (C). Unblocked session-by-session analyses revealed that NAc-, DMS-, and DLS-probe-implanted mice did not differ in their accuracy of VMCL test and all subjects became asymptotic –defined as no difference between consecutive mean percent correct scores –in the last 4 sessions (D). Similarly, groups did not differ significantly in the number of correction trials required (E), or latencies to make correct (F) or incorrect (G) choices. Interestingly, DMS-probe-implanted mice took slightly longer to collect their rewards (H), irrespective of session block. In our experience, however, this latency remains within the normal range for touchscreen tasks, which is usually around 1-2 seconds. Data presented as Mean + SEM, One- or Two-way RM ANOVA, ** p<0.01.

DAergic transmission within the striatum plays a crucial role in modulating corticostriatal plasticity (Centonze et al. 1999, 2003; Kerr and Wickens 2001), potentiating striatal NMDA function (Hallett et al. 2006), and exerting profound effects on coordinated activity of neurons in corticostriatal ensembles (Costa et al. 2006), but is thought to display regional hetereogeneity (Collins and Saunders 2020). To compare DAergic transmission across the striatum in learning and the expression of learning, we combined the recently developed GRABDA2m biosensor with fiber photometry recordings to characterize NAc-, DMS-, and DLS-DA dynamics in freely moving wild-type C57BL/6J mice during learning and expression of learning during the VMCL task. This setup has been described in detail elsewhere (Kljakic et al. 2022b; Skirzewski et al. 2022; Piantadosi et al. 2025) and enables recording of fast, phasic dynamics with high temporal and spatial specificity (Sun et al. 2020). Using this approach, we obtained robust and reliable recordings of extracellular DA dynamics across the NAc, DMS and DLS. For each region, we verified validated the sensor sensitivity with injections of systemic cocaine (10mg/kg, *i.p.*) (Fig. 1C, 1F, 1I) and the expression of GRABDA2M and fiber optic probe location (Supplementary Fig. 1D, 1G, 1J).

We examined DA dynamics during four key task events within trials of the VMCL task: (1) *trial initiation*, the point at which a mouse enters the reward tray to initiate a new trial, (2) *correct choice*, the point at which the correct right- or left-flanking position is nose-poked, (3) *incorrect choice*, the point at which the incorrect right- or left-flanking position is nose-poked, and (4) *reward collection*, the point at which the mouse collects its strawberry milkshake reward from the reward magazine. These task events are *‘animal-controlled’* as they are led by the subject, rather than by the investigator (Piantadosi et al. 2025). Although this flexibility provides subjects with greater control over the pacing and sequencing of behavior, it means that the timing of task events will vary across individual mice. Thus, to correlate behavior with neural signatures across groups of mice, we normalized and aligned our trial-based fiber photometry recordings from each mouse to distinct event windows (e.g. 10s prior to and following each correct choice event, with ‘time = 0’ as indicated by the dotted line on the X axis). This method allowed us to calculate the average DA signal for each subject during critical VMCL task events, irrespective of variations in timing across the session. The resulting data were then represented as heat maps to illustrate DAergic transients across trials of the same task event (Fig. 3D-F, J-L; Fig. 4D-F, J-L) or aggregated across all subjects within their respective probe-implanted groups and plotted as line graphs to visualize group differences (Fig. 3–4). In this way, although the event windows often overlapped (e.g. correct choice precedes reward collection by ∼1s), the neural signatures were time-locked at ‘time = 0’ to each event point with millisecond precision.

**Fig. 3.**
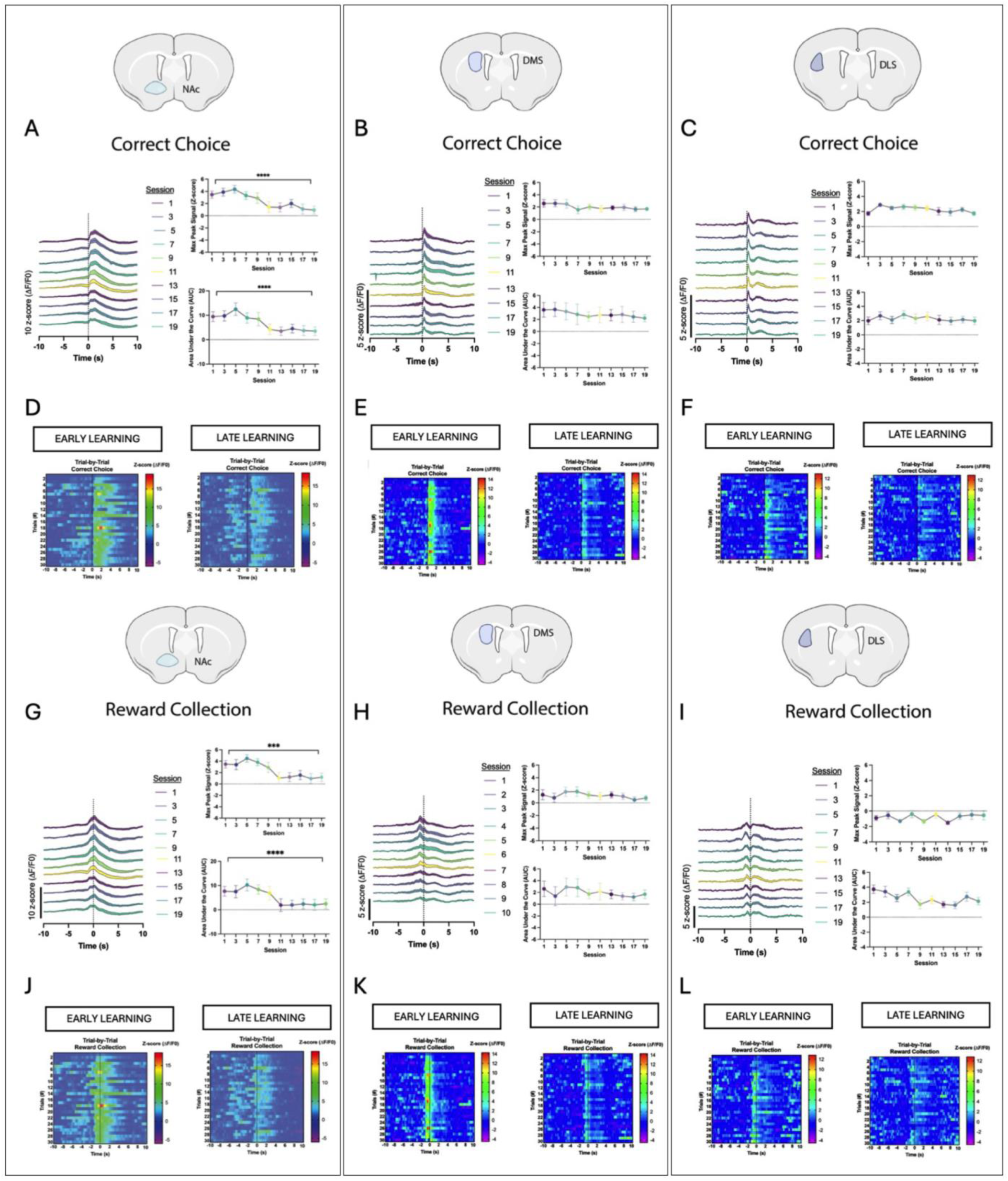
Striatal dopamine is tightly coupled to correct choice and reward collection, and is dynamic across visuomotor conditional learning in the NAc, while in the DMS and DLS it peaks at correct choice and remains stable throughout learning. (A-C) Normalized mean DA signal in the NAc, DMS, and DLS at correct choice across learning (every second session is plotted), along with their associated max peak and area under the curve quantifications. (D-F) Trial-by-trial heatmaps of one representative subject for NAc, DMS, and DLS at correct choice in early and late learning. (G-I) Normalized mean DA signal in the NAc, DMS, and DLS at reward collection across learning (every second session is plotted), along with their associated max peak and area under the curve quantifications. (J-L) Trial-by-trial heatmaps of one representative subject for NAc, DMS, and DLS at reward collection in early and late learning. Data presented as Mean + SEM, One-way RM ANOVA, *** p <0.001, **** p < 0.0001.

**Fig. 4.**
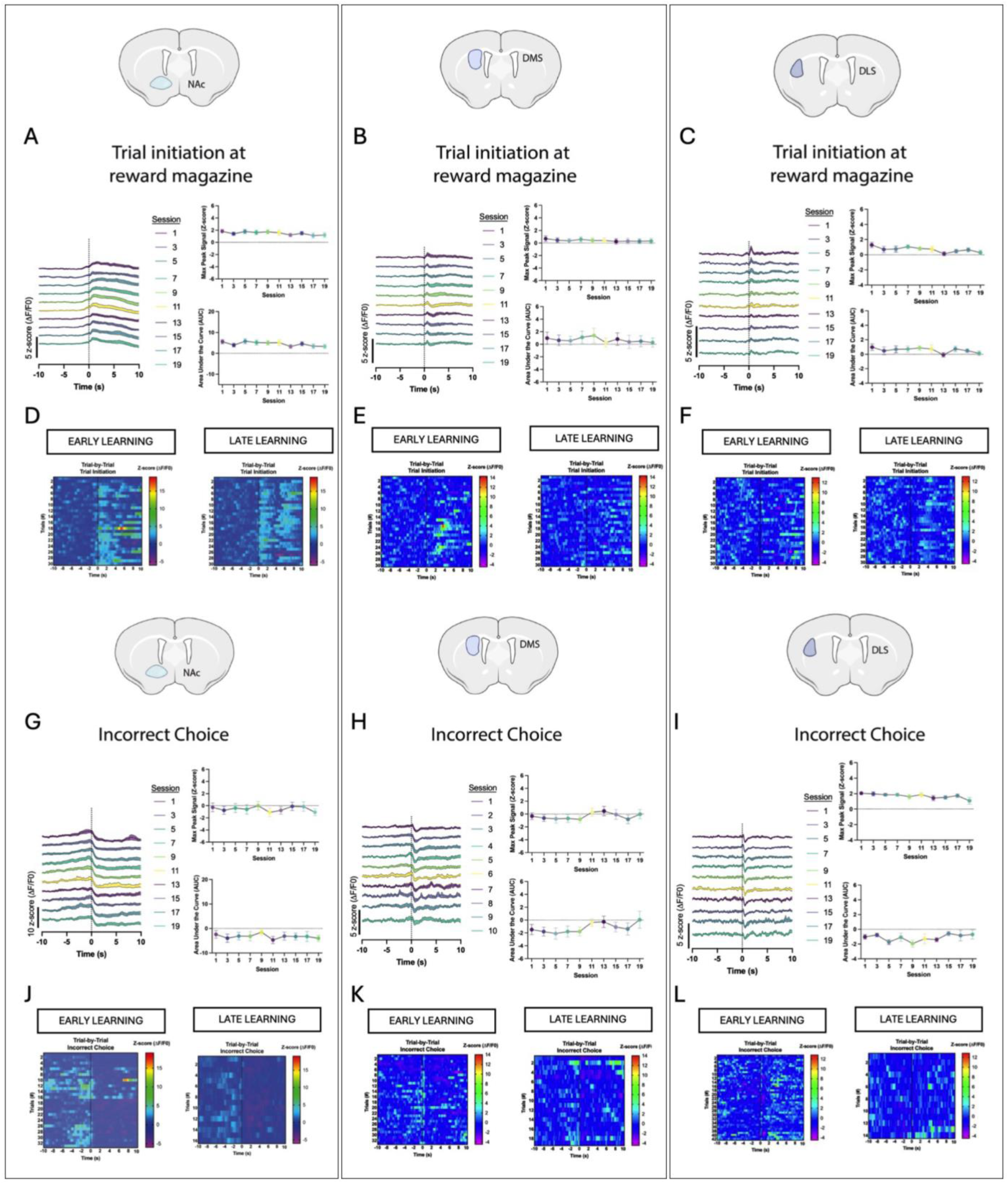
Striatal dopamine is tightly coupled to trial initiation and incorrect choice, and remains stable during visuomotor conditional learning. (A-C) Normalized mean DA signal in the NAc, DMS, and DLS at trial initiation at reward magazine across learning (every second session is plotted), along with their associated max peak and area under the curve quantifications. (D-F) Trial-by-trial heatmaps of one representative subject for NAc, DMS, and DLS at trial initiation at reward magazine in early and late learning. (G-I) Normalized mean DA signal in the NAc, DMS, and DLS at incorrect choice across learning (every second session is plotted), along with their associated max peak and area under the curve quantifications. (J-L) Trial-by-trial heatmaps of one representative subject for NAc, DMS, and DLS at incorrect choice in early and late learning.

Employing this method, we obtained robust evidence that NAc DA dynamics are tightly coupled to VMCL task events in early learning, with modest increases at trial initiation, but elevated and long-lasting responses triggered at correct choice that peak in amplitude at reward collection (Supplementary Fig. 1B, top panel). In contrast, NAc DA dipped in response to incorrect choice, although it did not descend significantly below baseline (Supplementary Fig. 2C; 1-way ANOVA --F3.868 = p<0.0001; Sidak’s *post-hoc* test: baseline vs correct choice (p<0.001), baseline vs trial initiation (p<0.001) and baseline and reward collection (p<0.01). The peak heights of these transients were significantly different from one another (Supplementary Fig. 2E; 1-way ANOVA, F3.736, p<0.001. Sidak’s *post-hoc* test: trial initiation vs incorrect choice (p<0.05), correct vs incorrect choice and incorrect vs reward collection (p<0.01)) and were highly robust, occurring on almost every trial (Fig. 3D-F, J-L; Fig. 4D-F, J-L (left panels)). Interestingly, these neural signatures differed greatly in their stability across learning (Supplementary Fig. 1B, bottom panel, Fig. 3A-C, G-I; Fig. 4A-C, G-I). DA transients for rewarding events, including correct choice and reward collection, for example, decreased dramatically across learning, supporting a reward prediction error (RPE)-like blunting of DA as the value of the prediction decreases across sessions (Schultz 2016). This was evidenced by a 2.5-3x fold reduction in max peak signal and area under the curve (AUC) for correct choice (Fig. 3A; Max Peak: 1-way ANOVA --F4.010, 44.11= 10.34, p<0.0001; AUC: 1-way ANOVA --F4.115, 45.27= 12.91, p<0.0001) and reward collection (Fig. 3G; Max Peak: 1-way ANOVA --F3.574, 39.31= 7.995, p<0.0001; AUC: 1-way ANOVA --F4.152, 45.67 = 8.742, p<0.0001). All statistical details are listed in Supplementary Table 1-2. Critically, this reduction was not present for negatively valence or neutral (Fig. 4A, G) events, ruling out the possibility that reductions in signal might be due to sensor- or fiber optic-related artefacts (e.g., photobleaching, fiber probe damage etc.).

Ramp-like activity was also evident in the NAc, as previously described (Hamid et al. 2015; Collins et al. 2016a; Mohebi et al. 2019; Collins and Saunders 2020; Kim et al. 2020; Farrell et al. 2022). Gradual increases in DA release have been reported as animals navigate their environment in pursuit of a distal reward (Howe et al. 2013) and during the completion of instrumental action sequences (Hamid et al. 2015; Collins et al. 2016b), likely providing motivational drive. Here, we found prolonged elevations in max peak signals in the NAc during early and late learning as subjects neared the choice period, for both correct (Supplementary Fig. 3B-C; paired t-test: p<0.01) and incorrect choices (Supplementary Fig. 3D-E; paired t-test: p<0.05).

Robust DA dynamics were also present in the DMS and DLS with several key distinctions. For example, DA dynamics in these dorsal regions were still highly coupled to VMCL task events early in learning (Supplementary Fig. 1E, H), had max peak signals that differed significantly from one another and were present on almost every trial (DMS: Supplementary Fig. 2I; 1-way ANOVA --F_3.803_ = p<0.01, Sidak’s *post-hoc* test: correct vs incorrect choice (p<0.01), correct choice vs reward collection (p<0.05). DLS: Supplementary Fig. 2M; 1-way ANOVA --F_1.382_ = p<0.0001, Sidak’s *post-hoc* test: baseline vs correct choice (p<0.001), baseline vs incorrect choice (p<0.05), baseline vs reward collection (p<0.0001); late learning: Supplementary Fig. 2O, 1-way ANOVA --F_0.7779_ = p<0.05, Sidak’s *post-hoc* test: trial initiation vs correct choice (p<0.001)). However, these dynamics were significantly more transient. For example, unlike NAc-DA which peaked around reward collection, DMS- and DLS-DA both peaked ∼200-400ms after correct choice, which occurred ∼500ms *prior* to reward collection (Supplementary Fig. 1E, H). This was reflected in a significant difference between the AUC following a correct choice in the NAc compared to the DLS and DMS (Fig. 5B; 1-way ANOVA --F_6.705_ = p<0.01, Sidak’s *post-hoc* test: NAc vs DMS (p<0.05), NAc vs DLS (p<0.01)). This finding, combined with the reward-related activity in the NAc, might indicate a role for the learning versus the expression of learning in the NAc and DMS/DLS, respectively.

**Fig. 5.**
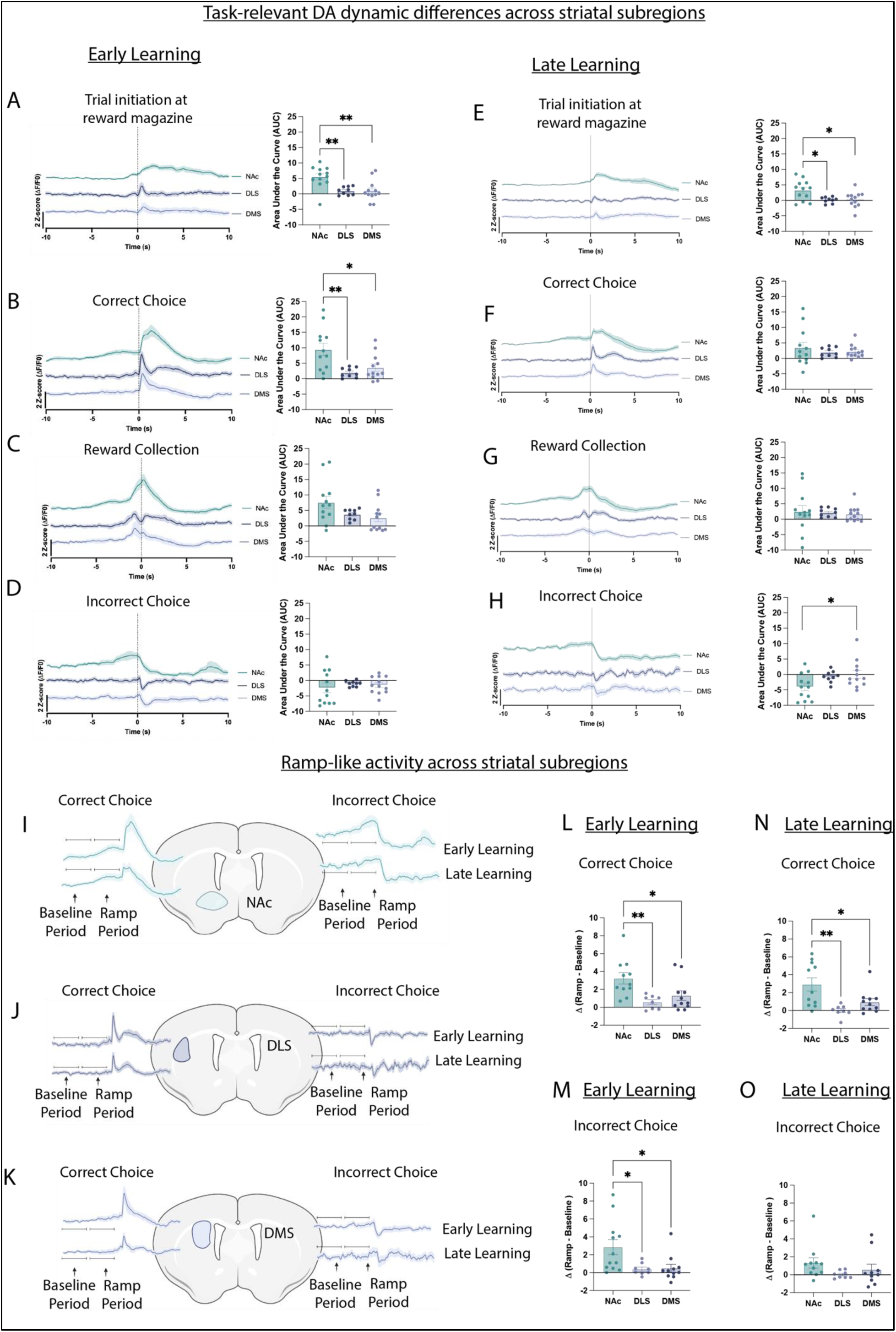
*in vivo* dopamine dynamics are longer-lasting and display more ramp-like activity in the nucleus accumbens than in dorsomedial or dorsolateral striatum. (A) Normalized mean dopamine (DA) signals across subjects with recordings in the nucleus accumbens (NAc), dorsolateral striatum (DLS) and dorsomedial striatum (DMS) were compared in early (Session 1, A-D) and late learning (Session 19, E-H) aligned to four key task events: Trial Initiation, Correct Choice, Reward Collection, and Incorrect Choice. Differences between groups was quantified by comparing the area under the curve (AUC) following each event. Ramp-like activity preceding choice points was quantified by comparing the AUC during a 5 second ‘baseline period’ and the 5 seconds ‘ramp period’ leading up to correct or incorrect choices, in the NAc (I), DLS (J) and DMS (K). Differences between groups were quantified by computing a differential between the ramp and baseline period following correct (L) and incorrect choices (N) in early learning (Session 1), and correct (M) and incorrect choices (O) in late learning (Session 19). Data presented as Mean + SEM, 1-way ANOVA, * p<0.05, ** p<0.01.

A significant difference in AUC among the three different regions was also found for trial initiation (Fig. 5A; 1-way ANOVA --F_8.722_ = p<0.01, Sidak’s *post-hoc* test: NAc vs DMS and NAc vs DLS (p<0.01)), but no difference was observed following reward collection or incorrect choice (Fig. 5C-D). These temporal differences fit well with other recent comparative studies *in vitro* (Calipari et al. 2012; Salinas et al. 2023a) and *in vivo* (Van Elzelingen et al. 2022; Salinas et al. 2023b; Eshel et al. 2023). Interestingly, the pattern of response at reward collection was also visually distinct between the DMS and DLS, as detected elsewhere (Seiler et al. 2022b; Eshel et al. 2023). Whereas the DMS showed minimal positive peak at reward collection (Fig. 3H), the DLS revealed a second (albeit smaller) positive peak ∼750 milliseconds after reward collection which lasted ∼3000ms (Fig. 3I).

Furthermore, unlike NAc-DA, DMS- and DLS-DA did not significantly decrease across learning in response to rewarding events (Supplementary Fig. 1E, H; Fig. 3B-C, H-I). For example, although there was numerically a mild (∼1.5X) reduction in max peak signal and AUC between the first and last session at correct choice, this reduction did not reach statistical significance (Fig. 3B). This lack of effect also held true for DMS-DA in response to reward collection, as well as for both correct choice and reward collection for DLS-DA (Fig. 3B, H). As a result, no significant differences were found between groups in the AUC following correct choice or reward collection in late learning (Fig. 5F-G). Furthermore, DA dynamics were stable for negatively valence and neutral events (Fig. 4B-C, H-I). Taken together with the dynamic reduction present in the NAc, this finding may provide further support for distinct roles of the NAc, and DMS/DLS in learning versus expression of learning, respectively.

Finally, although ramp-like activity preceded both correct and incorrect choice in NAc-probe-implanted mice, in DMS-probe-implanted mice ramp-like activity was present only prior to correct choices, and no ramp-like activity was found within the DLS (Supplementary Fig. 3). Correspondingly, comparisons between groups revealed a significantly greater differential between baseline and ramp periods in the NAc when compared to the DMS or DLS prior to correct choices in early (Fig 5L, 1-way ANOVA --F_6.312_ = p<0.01, Sidak’s *post-hoc* test: NAc vs DLS (p<0.01), NAc vs DMS (p<0.05)) and late learning (Fig 5N, 1-way ANOVA --F_5.135_ = p<0.05, Sidak’s *post-hoc* test: NAc vs DLS (p<0.05)). A significantly greater differential between the NAc and DLS was also found prior to making incorrect choices in early learning (Fig. 5M, 1-way ANOVA --F_7.378_ = p<0.01, Sidak’s *post-hoc* test: NAc vs DLS (p<0.01), NAc vs DMS (p<0.05)) but not late learning (Fig. 5O). Taken together, these experiments revealed clear and robust neural signatures across the striatum that tracked task events. Bidirectional DA dynamics were apparent at choice in the NAc, DLS and DMS, with each area exhibiting unique temporal dynamics.

### Probability probes reveal striatal dopamine dynamics reflect reward delivery and reward omission, irrespective of choice

The bidirectional DA responses observed near choice-point could reflect the animals’ choice, or the associated reward delivery or reward omission that follows. To disentangle these possibilities, we ran a subset of well-trained subjects (NAc: *N* =7, *n*=4^♂^, *n*=3^♀^); DMS (*N*=8, *n*=3^♂^, *n*=5^♀^); DLS (*N*=6, *n*=2^♂^, *n*=4^♀^) on a series of behavioral probes in which the probability of reward was manipulated (Fig. 6).

**Fig. 6.**
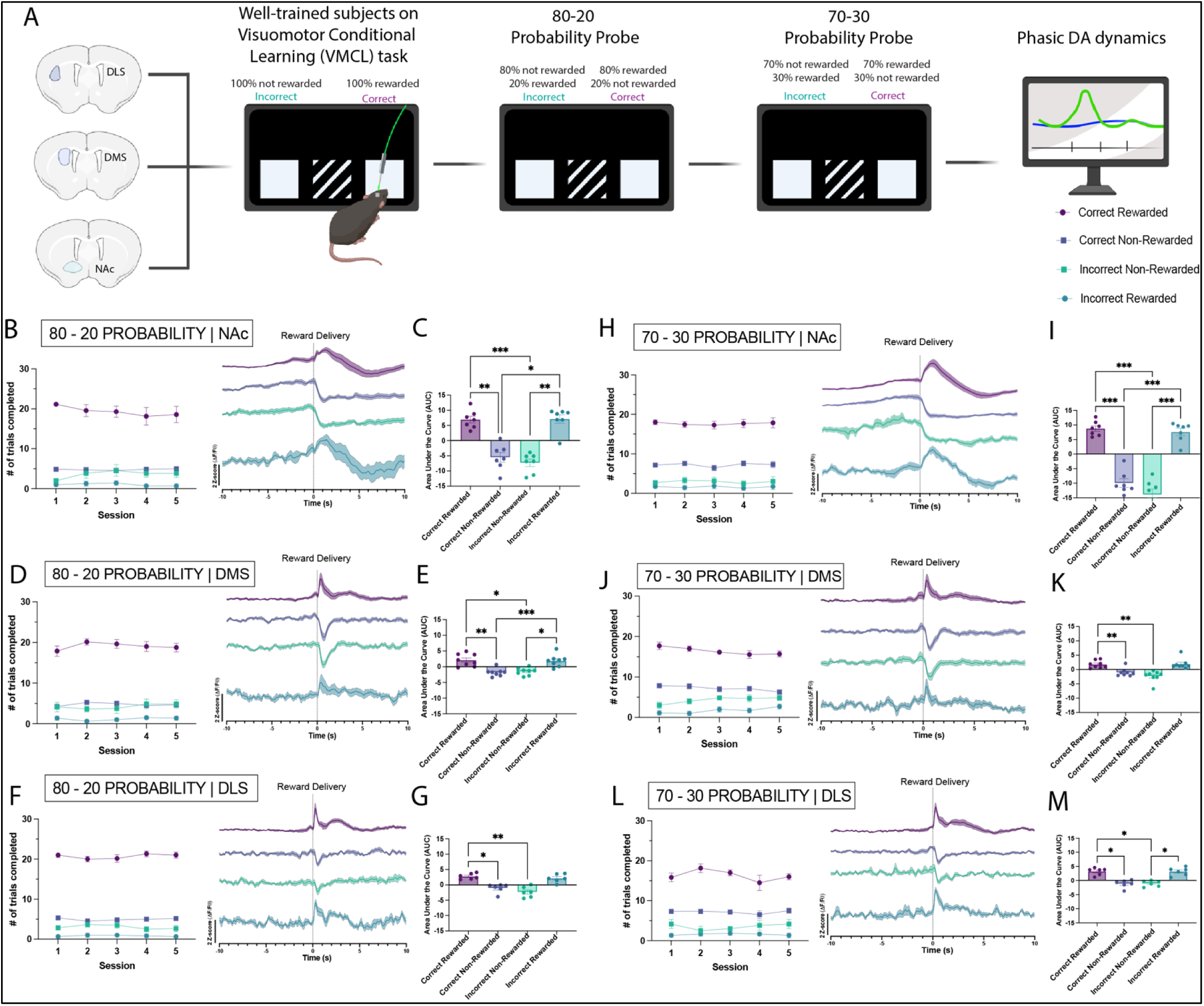
Manipulating reward probability reveals striatal dopamine tracks reward delivery and omission, irrespective of choice. (A) Following the completion of the Visuomotor Conditional Learning task, a subset of well-trained C57BL6J mice were tested on probability probe tests. In the 80-20 Probability Probe (A-G), the probability of gaining a reward following a correct choice was set to 80%, and the probability of gaining a reward following an incorrect choice was set to 20%. All subjects in the nucleus accumbens (NAc), dorsomedial striatum (DMS) and dorsolateral striatum (DLS) showed minimal behavioural adaptation across 5 sessions, and mean dopamine (DA) dynamics reflected reward delivery and reward omission, irrespective of choice (B-F). Quantifications of these patterns of activity were analyzed by calculating the area under the curve (AUC) following each trial condition (e.g. correct rewarded, correct non-rewarded, incorrect rewarded and incorrect non-rewarded) (C-G). In the 70-30 Probability Probe (A, H-M), the probability of gaining a reward following a correct or an incorrect choice shifted again, such as that 70% of correct choices were rewarded, and 30% of incorrect choices were rewarded. Similarly, all subjects in the NAc, DMS, and DLS displayed minimal behavioural adaptions across 5 sessions, and mean DA dynamics reflected reward delivery and reward omission, irrespective of choice (H-L). Quantifications of these patterns of activity were analyzed by calculating the AUC following each trial condition. Data presented as Mean + SEM, One-way ANOVA, *p<0.05, **p<0.01, ***p<0.001.

First, we ran an 80-20 probability probe, in which the probability of receiving a reward after a correct choice was set to 80 % (rather than 100%), and after an incorrect choice was set to 20% (rather than 0%). This provided us with 4 different trial conditions at choice: (a) *correct rewarded trials*, (b) *correct not rewarded*, (c) *incorrect rewarded trials* and (d) *incorrect not rewarded trials*. Across 5 consecutive sessions, performance was stable in all groups (Fig. 6B,D,F, 2-way RM ANOVA –NAc: Main effect of group, F_3,24_=95.70, p<0.0001); DMS: Main effect of group, F_3,28_=195.5, p<0.0001; DLS: Main effect of group, F_1.164,5.822_=626.8, p<0.0001). In fact, all subjects maintained a percent correct accuracy of >85% across all 5 sessions. We therefore time-locked our DA dynamics to the choice period and visualized the average response in our four choice conditions.

These recordings revealed clear, unequivocal evidence that DA dynamics across the striatum reflect reward delivery and reward omission, and not choice, *per se*. For example, when a correct choice was made, DA dynamics across all three regions demonstrated phasic elevations only when the trial was rewarded (*correct rewarded trials*). When a correct choice was made that was not rewarded (*correct non-rewarded trials*), we found a depression in signal that resembled that found on incorrect trials during VMCL testing. Correspondingly, when an incorrect choice occurred (although these were infrequent during probe trials), the expected depression in signalling occurred only on non-rewarded trials (*incorrect non-rewarded trials*). Incorrect choices that were rewarded (*incorrect rewarded trials*) were accompanied by DA dynamics across all regions that looked remarkably like the correct rewarded trials (although the error was much greater as these trial types were extremely low in number (∼1 trial per session per subject)). These differences were supported by significant differences in AUC between trial conditions in NAc-probe-implanted mice (Fig. 6C, 1-way ANOVA --F_35.38_ = p<0.0001, Sidak’s *post-hoc* test: correct rewarded vs correct non-rewarded trials and incorrect rewarded vs incorrect non-rewarded trials (p<0.01), correct rewarded vs incorrect non-rewarded trials (p<0.001), correct non-rewarded vs incorrect reward trials (p<0.05)), DMS-probe implanted mice (Fig. 6E, 1-way ANOVA --F_20.07_ = p<0.0001, Sidak’s *post-hoc* test: correct rewarded vs correct non-rewarded trials (p<0.01), correct rewarded vs incorrect non-rewarded trials and incorrect rewarded vs incorrect non-rewarded trials (p<0.05), correct non-rewarded vs incorrect reward trials (adj. p<0.001)) and DLS-probe implanted mice (Fig. 6G, 1-way ANOVA --F_16.29_ = p<0.001, Sidak’s *post-hoc* test: correct rewarded vs correct non-rewarded trials (p<0.05), correct rewarded vs incorrect non-rewarded trials (p<0.01)).

To further probe the behavioral flexibility of these mice, we subsequently reduced the probability of reward to 70-30 and found same pattern of behavior. No behavioural changes were observed across the 5 consecutive sessions (Fig. 6H,J,L, 2-way RM ANOVA –NAc: Main effect of group, F_3,25_=177.9, p<0.0001; DMS: Main effect of group, F_3,24_=309.9, p<0.0001; DLS: Main effect of group, F_3,24_=95.70, p<0.0001), and once again the DA dynamics in all regions reflected reward delivery and omission, and not choice (Fig. 6I,K,M, 1-way ANOVA –NAc: F_61.40_ = p<0.0001, Sidak’s *post-hoc* test: correct rewarded vs correct non-rewarded trials, correct rewarded vs incorrect non-rewarded trials, correct non-rewarded vs incorrect rewarded and incorrect rewarded vs incorrect non-rewarded trials (p<0.001). DMS: F_11.42_, p<0.01, Sidak’s *post-hoc* test: correct rewarded vs correct non-rewarded trials and correct rewarded vs incorrect non-rewarded trials (p<0.01). DLS: F_22.23_= p<0.01. Sidak’s *post-hoc* test: correct rewarded vs correct non-rewarded trials, correct rewarded vs incorrect non-rewarded trials, correct non-rewarded vs incorrect rewarded, and incorrect rewarded vs incorrect non-rewarded trials (p<0.05). Taken together, this work unequivocally confirms that DA transients in the NAc, DMS and DLS in our task tracks reward delivery and reward omission, irrespective of choice.

### DLS-projecting nigrostriatal but not NAc-projecting mesolimbic dopaminergic afferents are necessary for learning

Correlation does not imply causation. While we observed clear and dynamic DAergic neuromodulation across the striatum, it remained unclear whether these signals were strictly necessary for the VMCL task. We therefore tested whether chemogenetic inhibition of DA transmission to the striatum via either the nigrostriatal or mesolimbic DA pathways impairs learning and/or the expression of learning. When combined with the synthetic agonist CNO, Designer Receptors Exclusively Activated by Designer Drugs (DREADDs) have been shown to reduce phasic firing of neurons by inducing membrane hyperpolarization through reduced cyclic adenosine monophosphate (cAMP) signaling and activation of G protein-coupled inwardly rectifying potassium (GIRK) channels (Armbruster et al. 2007; Stachniak et al. 2014) and is sensitive to DAergic pathway manipulations *in vivo* (Runegaard et al. 2018, 2019; Krishnan et al. 2022; Chen et al. 2022). Thus, we used the inhibitory DREADD hM4Di to assess how transient reductions of nigrostriatal or mesolimbic DA via inhibitory GPCR signaling affects behavior.

Specifically, we injected a retrograde AAV expressing Cre-dependent hM4D(Gi) (AAV-hSyn-DIO-hM4Di(Gi)-mCherry) (Krashes et al. 2011) bilaterally into either the DLS or NAc of DAT-IRES-Cre mice (Fig. 7A). This enabled selective expression of hM4D(Gi) in either the nigrostriatal DA pathway projecting to the DLS or the mesolimbic DA pathway projecting to the NAc, respectively. Hereafter, these groups were called the *DLS-projecting nigrostriatal DA inhibition* (*N*=15, *n*=8^♂^, *n*=7^♀^) and the *NAc-projecting mesolimbic DA inhibition* (*N*=17, *n*=7^♂^, *n*=10^♀^) groups. To control for off-target effects of CNO (MacLaren et al. 2016; Gomez et al. 2017) and to act as our between-subjects control group, we also injected a retrograde control transgene (rAAV-hSyn-Dio-mCherry) into the DLS and NAc of DAT-IRES-Cre mice, hereafter called our *nigrostriatal DA control* (*N*=18, *n*=9^♂^, *n*=9^♀^) group and *mesolimbic DA control* group (*N*=19, *n*=8^♂^, *n*=11^♀^), respectively.

**Fig. 7.**
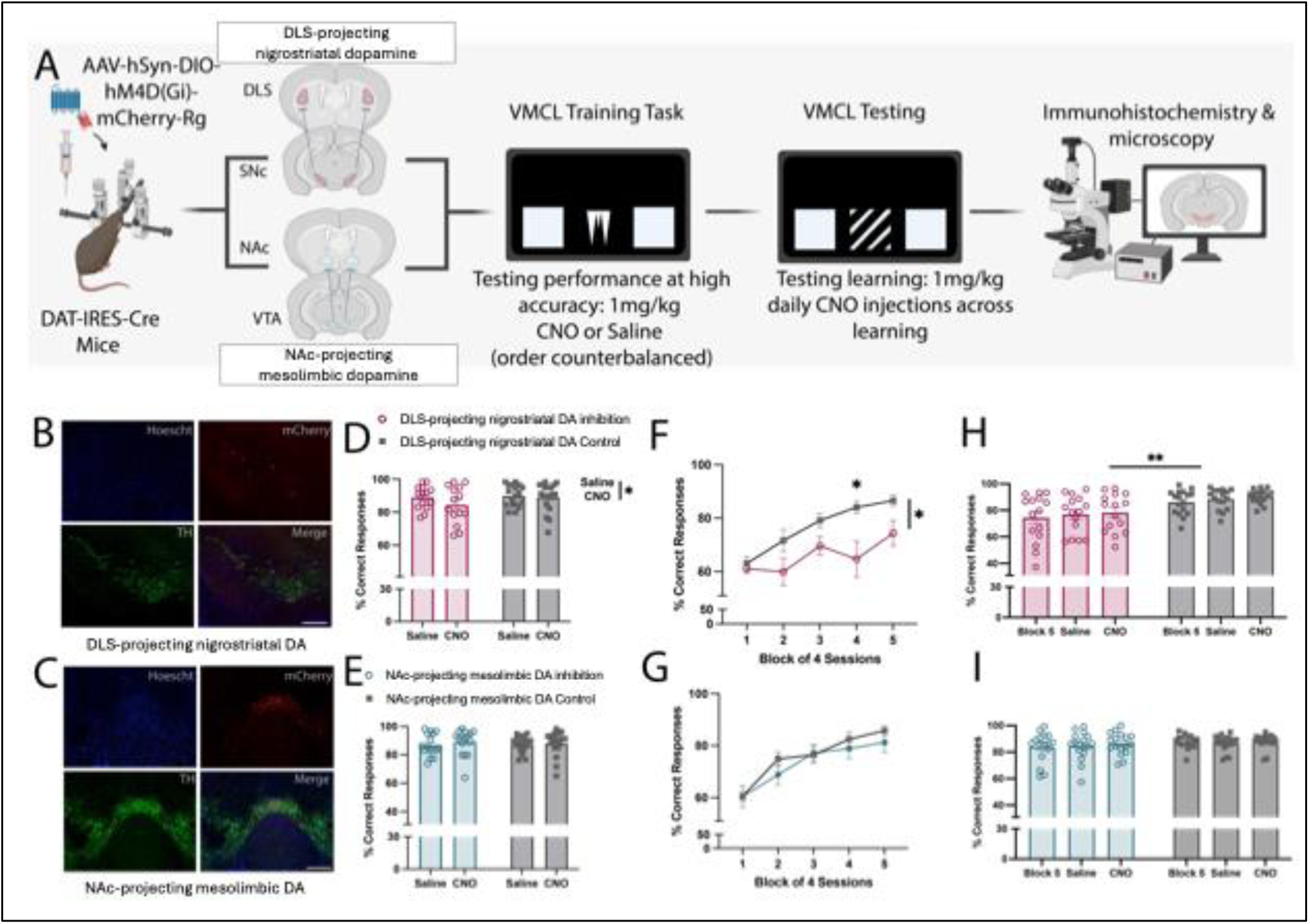
DLS-projecting nigrostriatal dopamine but not NAc-projecting mesolimbic dopamine is necessary for visuomotor conditional learning. (A) DAT-IRES-Cre mice underwent stereotaxic surgery for viral infusion of the retrograde inhibitory DREADD hM4D(Gi) or an empty AAV vector control into either the dorsolateral striatum (DLS) (which retroactively targeted the nigrostriatal dopamine (DA) pathway) or the nucleus accumbens (NAc) (which retroactively targeted the mesolimbic DA pathway). Subsequently, mice were trained on the Visuomotor Conditional Learning (VMCL) training task until they reached high accuracy, and then received acute injections of saline or the DREADD agonist clozapine N-Oxide (CNO) (1mg/kg, i.p.). This enabled the assessment of whether DLS-projecting nigrostriatal and NAc-projecting mesolimbic DA is necessary for the expression of previous learning. Next, subjects moved onto the VMCL test for 28 sessions, where they received 20 daily CNO treatments (1mg/kg, i.p.), followed by 4 sessions of saline relief and 4 sessions of CNO reinstatement. This enabled assessment of DLS-projecting nigrostriatal and NAc-projecting mesolimbic DA in new learning and the impact of temporary inhibition relief and reinstatement. Representative coronal 40x magnification brain sections showing colocalization of the mCherry-tagged viral vector and DA neurons, targeted with tyrosine hydroxylase (TH) (mCherry: red; TH: green; Hoescht: blue) in nigrostriatal substantia nigra pars compacta (SNc) neurons (B) and mesolimbic ventral tegmental area (VTA) neurons, scale bar 500µm (C). Relative to controls, subjects with nigrostriatal DA inhibition were not impaired in the expression of learning (D) but displayed significant impairments in acquiring new contingencies (F). This impairment was maintained throughout saline relief and CNO reinstatement (H). In contrast, mesolimbic inhibition had no impact on the expression of previous learning (E) nor the acquisition of new ones (G-I). Data presented as Mean + SEM, Two-way ANOVA or Two-way RM ANOVA, * p<0.05; ** p<0.01.

First, we verified that these stereotaxic injections into the striatum, targeting axon terminals, back-propagated to the midbrain neurons and induced expression of AAV-tagging mCherry, as previously reported (Chen et al. 2022). mCherry was detected with expression confined to tyrosine hydroxylase (TH)-positive neurons. DLS-injected groups (i.e. DLS-projecting nigrostriatal DA inhibition and nigrostriatal DA control) showed clear and selective expression in the lateral portions of the SNc (Fig. 7B, Supplementary Fig. 4B), with similar expression on control and CNO groups (Supplementary Fig. 4D, unpaired t-test, p=0.0455). NAc-injected groups (i.e. NAc-projecting mesolimbic DA inhibition and mesolimbic DA control) had equivalent selectivity of expression in the medial portions of the VTA (Fig. 7C, Supplementary Fig. 5A, D, unpaired t-test, p=0.0229).

Although these manipulations were selective to dopaminergic projections, knock-on effects of this dopaminergic depletion would be expected to be wide-ranging, impacting many aspects of striatal circuit function. Therefore, we performed functional c-fos analysis to assess broad striatal dysfunction. CNO was found to significantly lower neural activity in DLS, NAc, SN, and VTA (Supplementary Fig. 4C, unpaired t-test, p=0.0150). Supplementary Fig. 5A-C, unpaired t-test, p=0.0346). Taken together, these results support that our chemogenetic manipulations resulted in a partial functional inactivation of the DLS-projecting nigrostriatal and NAc-projecting mesolimbic pathways and that this inactivation was approximately equal in both pathways.

With this confirmed, we set out to explore the role of nigrostriatal and mesolimbic DA in the acquisition and expression of visuomotor conditional learning. Our experimental design drew inspiration from studies that distinguish between associative learning and its expression (Brain et al. 1993; Atallah et al. 2007). Although intimately linked, the acquisition and expression of previously learned actions has been suggested to rely on distinct striatal circuits within an actor-critic framework (Joel et al. 2002; O’Reilly and Frank 2006; Atallah et al. 2007). However, the specific roles of mesolimbic and nigrostriatal DA within these networks remains unknown.

We therefore exposed mice to a series of behavioral manipulations designed to evaluate the expression of previously acquired contingencies and the acquisition of novel ones. First, we trained subjects on the VMCL training task until they achieved a high level of accuracy, ensuring both experimental and control groups reached comparable proficiency before any chemogenetic interventions. No significant difference in sessions to reach a criterion of ≥77% accuracy over two consecutive sessions was found between the DLS-projecting nigrostriatal DA inhibition group and their controls (Supplementary Fig. 6B), nor between the NAc-projecting mesolimbic DA inhibition group and their controls (Supplementary Fig. 6C). Following this confirmation, we then compared VMCL performance following acute doses of systemic CNO (1mg/kg, *i.p*.) and saline, administered 30 minutes prior to task onset. The order in which CNO and saline was given was counterbalanced across animals. This allowed us to compare the influence of DLS-projecting nigrostriatal and NAc-projecting mesolimbic DA on the expression of previously acquired contingencies.

Unexpectedly, we found acute DLS-projecting nigrostriatal and NAc-projecting mesolimbic DA inhibition had little-to-no impact on expression of previously learning (Fig. 7D-E, 2-way RM ANOVA –Nigrostriatal DA inhibition: Main effect of drug, F_1,30_ = 4.414, p<0.05. Supplementary Fig. 6). Specifically, although there was a small but significant main effect of drug treatment (CNO vs Saline) for % correct responses in the nigrostriatal groups, which appeared to be driven by poorer performance in the DLS-projecting nigrostriatal DA inhibition group, there was no significant drug X group interaction (Fig. 7D). There was also no significant difference between the DLS-projecting nigrostriatal DA inhibition group and their controls on supplementary performance measures including perseveration error, missed or correction trials, and latencies to perform the task (Supplementary Fig. 6D,F,H,J,L,N). Similarly, mesolimbic DA inhibition had little effect on the expression of previous learning. We found no significant effect on % correct responses (Fig. 7D), perseveration error, correction trials or task latencies (Supplementary Fig. 6E,G,K,M,O).

Next, we assessed the influence of DLS-projecting nigrostriatal and NAc-projecting mesolimbic DA inhibition on learning. Here, we conducted a series of experiments spanning 28 consecutive sessions of the VMCL Test. The initial 20 sessions involved daily systemic injections of CNO (1mg/kg, *i.p*.), administered 30 minutes prior to task onset (Fig. 7F-G, Supplementary Fig. 7). Subsequently, we introduced 4 sessions of saline relief, followed by 4 sessions of CNO reinstatement (Fig 7H-I, Supplementary Fig. 8). This experimental design enabled us to assess the impact of DAergic inhibition throughout the learning phase and investigate the impact on the expression of learning when inhibition was temporarily relieved and then reinstated.

Inhibition during the VMCL test implicated DLS-projecting nigrostriatal but not NAc-projecting mesolimbic DA afferents in learning of the task. Relative to their controls, mice undergoing DLS-projecting nigrostriatal DA inhibition showed impaired % correct responses, resulting in a >15% reduction in their final accuracy in block 5, and a significant interaction with session (Fig. 7F, 2-way RM ANOVA --main effect of group: F_1, 31_= 6.977, p<0.05; main effect of session: F_2.876, 89.16_=28.97, p<0.0001; group x session interaction: F_4,124_=3.377, p<0.05. Sidak’s *post-hoc* test: block 4, p< 0.05). These mice also exhibited a higher perseveration index (Supplementary Fig. 7B, RM-Mixed Effects Model --main effect of group: F_1, 31_= 5.637, p<0.05; main effect of session: F_3.295, 93.09_= 10.40, p<0.0001) and took longer to make an incorrect choice (Supplementary Fig. 7L, RM-Mixed Effects Model --main effect of group: F_1,31_= 6.246, p<0.05; main effect of session: F_3.461,106.4_= 31.51, p<0.0001), both of which may suggest a decreased sensitivity to negative feedback. Furthermore, although their littermate controls reached asymptote in the last 6 sessions of the VMCL Test –defined as no difference between consecutive mean percent correct scores –subjects with DLS-projecting DA inhibition did not display evidence of stable asymptotic performance (Supplementary Fig. 7N, 2-way RM ANOVA --main effect of group: F_1, 31_= 8.002, p<0.01; main effect of session: F_6.178,190.2_= 9.338, p<0.0001; group x session interaction: F_19,585_= 2.384, p<0.001). However, no significant differences were found in the number of correction trials, percentage of missed trials or latencies to make a correct choice and collect rewards (Supplementary Fig. 7D,F,H,J).

We then tested animals’ performance when inhibition was temporarily relieved and then reinstated (Fig. 7H, Supplementary Fig. 8B,D,F,H,J,L). When comparing the average performance of mice undergoing DLS-projecting nigrostriatal DA inhibition and controls across the four sessions with CNO blockade in block 5, followed by 4 sessions on saline relief and then 4 sessions of CNO reinstatement, we found continuing impairments in VMCL performance, and no positive effects of saline relief. For example, DLS-projecting nigrostriatal DA inhibition caused enduring impairments on % correct responses (Fig. 7H, RM -Mixed Effects Model --main effect of drug: F_1.567,46.24_= 4.121, p<0.05; main effect of group: F_1,30_= 9.036, p<0.01). These mice also displayed elevated perseveration index (Supplementary Fig. 8B, RM-Mixed Effects Model --main effect of group: F_1,31_= 5.915, p<0.05; main effect of session: F_1.944,53.46_=3.482, p<0.05), correction trials (Supplementary Fig. 8D, RM-Mixed Effects Model --main effect of drug: F_1.678,48.68_= 4.300, p<0.05; main effect of group: F_1,31_= 11.68, p<0.01, missed trials (Supplementary Fig. 8F, RM-Mixed Effects Model --main effect of group: F_1,31_= 6.396, p<0.05; drug x group interaction: F_2,51_= 4.154, p<0.05. Sidak’s *post-hoc* test: nigrostriatal inhibition vs controls during CNO treatment, p<0.01), and latencies to make a correct (Supplementary Fig. 8J, 2-way RM ANOVA --main effect of group: F_1, 29_= 7.122, p<0.05) and incorrect choice (Supplementary Fig. 8L, 2-way RM ANOVA --main effect of group: F_1,30_= 9.790, p<0.01). Together, these data implicate DLS-projecting nigrostriatal DA in VMCL learning, with a pattern of behavior that is not ameliorated by inhibition relief or reinstatement.

In stark comparison, NAc-projecting mesolimbic DA blockade had little-to-no impact on VMCL learning (Fig. 7G, 2-way RM ANOVA --main effect of session: F_3.053,103.8_= 51.61, p<0.0001). No significant differences were found between the NAc-projecting mesolimbic DA inhibition and mesolimbic control group in accuracy (% correct responses), perseveration index, missed trials and task latencies (Fig. 7G, Supplementary Fig. 7C,E,G,I,L,N). Furthermore, both NAc-projecting mesolimbic DA inhibition and control groups reached stable asymptotic performance levels by the last 6 sessions, with no group differences (Supplementary Fig. 7O, RM-Mixed Effects Model --main effect of session: F_19, 646_= 19.16, p<0.0001). Correspondingly, no significant differences were found between groups during saline relief and CNO reinstatement (Supplementary Fig. 8C,E,G,I,K,M). Taken together, this work demonstrates that DLS-projecting nigrostriatal DA afferents, but not NAc-projecting mesolimbic DA ones, are necessary for learning the VMCL task.

## Discussion

DAergic transmission across the striatum plays a pivotal role in modulating corticostriatal pathways that underlie cognition, and dysfunction in these pathways is a defining characteristic of numerous neurodegenerative and neuropsychiatric disorders. S-R learning is a crucial associative process thought to drive automatic behaviors (Dickinson 1985; Holland 2008; Ostlund and Balleine 2008; Malvaez and Wassum 2018). In the present study we studied the role of striatal dopaminergic circuits in learning using the touchscreen VMCL task (Dickinson 1985), designed to assess S-R learning. First, using *in vivo* fiber photometry combined with a genetically encoded GRAB_DA2M_ DA biosensor, we monitored real-time fluctuations of extracellular DA during VMCL in three striatal subregions: the NAc, DMS and DLS. We identified distinct, phasic DA transients across all three regions that closely correlated with task events and could be interpreted within a reward prediction error (RPE) framework (Garris et al. 1994a; Schultz et al. 1997; Mohebi et al. 2024a). For example, all three regions exhibited bidirectional responses at choice points, characterized by elevated DA levels following a correct choice, and decreased levels following an incorrect choice. Notably, our probability probes revealed that these responses correlated with reward collection and reward omission, respectively. Rewarded trials, whether preceded by a correct (correct-rewarded) or incorrect (e.g. incorrect-rewarded) choice, correlated with DA increases, while non-rewarded trials, whether preceded by an incorrect (e.g. incorrect-non-rewarded) or correct (correct-non-rewarded) choice, correlated with DA decreases.

In addition to these similarities, we also found differences in DA dynamics between the different striatal subregions, particularly in the timing of peak responses, the presence of ramp-like activity, and stability across learning. For example, DA transients in the dorsal regions (DLS and DMS) exhibited consistently faster kinetics compared to the NAc, with maximum peak signals occurring immediately following a correct choice. In contrast, the ventral NAc exhibited a comparatively delayed peak, occurring during the subsequent reward collection period. These findings align well with kinetics recently observed in mice undergoing Pavlovian conditioning (Salinas et al. 2023a; Mohebi et al. 2024b) and instrumental behavior (Mohebi et al. 2024b), and the DA transients induced by electrical stimulation seen in brain slices and *in vivo* (Garris et al. 1994b; Calipari et al. 2012; Mohebi et al. 2024b). Furthermore, although ramp-like activity was not seen in the DLS, and only modest evidence was found in the DMS, robust ramping was observed in the NAc as mice neared choice points. These distinctions are well supported within the literature on VTA-NAc DA circuits (Hamid et al. 2015; Collins et al. 2016a; Mohebi et al. 2019; Collins and Saunders 2020; Kim et al. 2020; Farrell et al. 2022), in which ramping activity is often interpreted as tracking value, goal proximity or motivation. In contrast, little evidence for ramping has been found in the DLS (Hamid et al. 2021; Seiler et al. 2022b), although some evidence has been found in the DMS (Hamid et al. 2021; Seiler et al. 2022b). Finally, although the maximum peak signalling during rewarding events (e.g. correct choice and reward collection) remained relatively stable within the DLS and DMS as subjects acquired the VMCL task, transients decreased dramatically within the NAc across learning, as described previously (Schultz 2010, 2013).

These differences can be interpreted in several, potentially overlapping, ways. For example, these variations in phasic activity may result from differences within the local microenvironment. For instance, the dorsal striatum has been shown to possess greater uptake, D2-receptor mediated regulation, and density and phosphorylation of DA transporters than the NAc, collectively contributing to faster DA clearance (Rothblat and Schneider 1997; Reynolds et al. 2001; Gowrishankar et al. 2018; Nolan et al. 2020). Consequently, the swift and transient responses in the DLS and DMS may arise from quick clearance following DA peaks, whereas the prolonged responses in the NAc may reflect the DA response to several overlapping events. Moreover, these dynamics may indicate variations in the local modulation of DA release across the striatal subregions by interneuron populations, including cholinergic interneurons. Extensive research suggests that cholinergic interneurons play a crucial role in regulating striatal DA (Exley and Cragg 2008; Threlfell et al. 2010; Cachope et al. 2012; Kljakic et al. 2022c; Skirzewski et al. 2022), in part through nicotinic and muscarinic receptors (Mechawar; Jones et al. 2001; Zhou et al. 2003; Threlfell et al. 2012b). For example, muscarinic agonists have been found to potentiate DA peak transients and decay constants (Shin et al. 2015), and the distribution and function of muscarinic (Threlfell et al. 2012c) receptors are heterogeneous across the striatum. Specifically, the regulation of DA release is supported by M2 and M4 acetylcholine receptors in the dorsal striatum, but only M4 receptors in the ventral striatum (Threlfell et al. 2010). Whether this distinction can give rise to the distinct patterns of DAergic transmission reflected in our recordings remains unknown.

Finally, this variation in signalling during distinct behaviorally relevant events across the different striatal subregions may also reflect unique functions of these regions during learning (Keiflin and Janak 2015). DA-neuron burst-firing at the time of an action (e.g., correct choice) has previously been suggested to mediate aspects of decision-making or choice enactment, while DA responses during the delivery of reward has been proposed to facilitate learning (Olds and Milner 1954; Keiflin and Janak 2015; Hiebert et al. 2017). These functions may not be evenly distributed across the different striatal subregions. This interpretation may also be supported by the differences in the stability of DA dynamics we observed across the striatum during learning. The maximum peak signalling at correct choice in the DLS and DMS, for example, remained relatively stable across learning, supporting a constant role for these regions in action / choice enactment. In contrast, however, transients in the NAc peaked at reward collection but decreased significantly as subjects acquired the task, suggesting a role for NAc DA in early learning of the task (Horvitz 2009; Bermudez and Schultz 2014). As a more rigorous test, we ran probability probes and found that responses in all three regions correlated with the presence or absence of reward, and not with correct or incorrect choice. Thus, it was difficult to ascertain specific functional roles of DA in the three regions from the profile of DA transients alone. One area in which this has been extensively studied RPE, defined as the discrepancy between actual and predicted rewards (Schultz et al. 1997; Bayer and Glimcher 2005; Hart et al. 2014). While some studies show relatively conserved RPE-like responses across the striatum (Tsutsui-Kimura et al. 2020; Van Elzelingen et al. 2022), others have identified functional heterogeneity of DA release patterns across the striatum that do not fit easily within this RPE framework (Howe and Dombeck 2016; Menegas et al. 2017; Engelhard et al. 2019; van Elzelingen et al. 2022). These findings challenge the notion that DA uniformly encodes RPEs across the striatum and instead suggests that dopaminergic signaling may be tailored to the computational demands of specific striatal subregions. Therefore, to test the functional necessity of striatal DA for VMCL learning, we carried out causal experiments using chemogenetic inhibition of DLS-projecting nigrostriatal afferents and NAc-projecting mesolimbic afferents during learning of the VMCL task. We found that inhibiting DA release from nigrostriatal or mesolimbic pathways during had minimal impact on the expression of previous learning. However, inhibiting these pathways during learning resulted in a dissociation between regions: selective effects were observed after DLS-projecting nigrostriatal DA inhibition, but not after NAc-projecting mesolimbic DA inhibition. These findings raise several intriguing points for discussion.

Firstly, the absence of impairment following DLS-projecting nigrostriatal DA inhibition during the expression of previous learning, in contrast to the pronounced impairment observed during the new learning, is consistent with previous pharmacological observations of reduced responsiveness to DA antagonism with extended training of appetitive behaviors (Beninger and Hahn 1983; Horvitz and Ettenberg 1991; Choi et al. 2005). This diminished responsiveness has been shown to be mediated by D1 receptor activity rather than D2 receptor activity (Choi et al. 2009) and may signify a shift of reliance from DA-modulated corticostriatal-basal ganglia circuits to direct corticocortical mediation (Houk and Wise 1995; Carelli et al. 1997; Ashby et al. 2007), or a decreasing role for DAergic modulation of intra-striatal glutamatergic circuits (Horvitz 2002; O’Donnell 2003). It must be noted, however, that our chemogenetic strategy targeted only a subset of nigrostriatal TH+ cells. Indeed, in our mice DREADD expression appeared to be relatively low. It is notable that the inhibition of an only subset of dopaminergic SN-DLS neurons led to a significant deactivation of the DLS and pronounced behavioural effects, suggesting that the specific behaviour we studied may be particularly sensitive to disruption of nigrostriatal dopamine. Nevertheless, it is conceivable that the subtotal inactivation achieved was not enough to disrupt behaviours that were preserved in the face of this inactivation, for example the expression of firmly established learning. Moreover, our viral chemogenetic strategy targeted DLS-projecting nigrostriatal dopamine neurons, and so such preserved behaviours may have been supported by functionally intact DMS-projecting dopamine neurons.

Secondly, absence of behavioral consequences resulting from mesolimbic DA inhibition in either the learning or the expression of learning is notable, particularly considering the dynamic DA transients observed in the NAc throughout the task. However, this is not the first report of neural correlates failing to align with behavioral necessity. Numerous instances exist where task-correlated neural signatures were found to be functionally unnecessary (Tremblay et al.; Schmaltz and Theios 1972; Solomon and Moore 1975; Berger and Thompson 1978; Darvas et al. 2014a; Fraser and Janak 2017a; Musall et al. 2019; Salinas et al. 2023a). These data also align with the growing body of literature supporting the necessity of nigrostriatal DA, as opposed to mesolimbic DA, for instrumental learning (for detailed discussion, refer to (Yin et al. 2008a; Salamone and Correa 2012a). For instance, studies have demonstrated that DA-depleting lesions using 6-OHDA or DA receptor blockade with α-flupenthixol induced profound and long-lasting deficits in conditional discrimination tasks when injected into the dorsal striatum (which affects nigrostriatal DA), but not when administered into the NAc (which affects mesolimbic DA)(Robbins et al. 1990a). In other studies, genetic depletion of DA in the nigrostriatal pathway impaired instrumental action, whereas depletion in the mesolimbic pathway did not (Szczypka et al. 2001a; Sotak et al. 2005; Robinson et al. 2007). Taken together with our current findings, these studies support the notion that nigrostriatal DA may be more essential than mesolimbic DA for habit learning.

This conclusion may seem at odds with that arrived at by others using somewhat different methodology (Atallah et al. 2007; Hiebert et al. 2017, 2019). For example, in a study using rats trained on a two-alternative forced choice task (A+ v. B-) with pharmacological inactivation of either the ventral or dorsal striatum, a dissociable role in learning and the expression of learning, respectively, was identified (Atallah et al. 2007). However, this discrepancy may be attributed, at least in part, to variations in experimental designs and methods. Our study focused on behavior designed to be contingent on S-R associations, whereas the two-alternative forced choice task used in the Atallah et al. (2007) study probably involves multiple associative processes. Furthermore, while we investigated the role of striatal DA, Atallah et al. (2007) used pharmacological inactivation of the dorsal or ventral striatum with the GABA-A receptor agonist muscimol and the NMDA receptor antagonist AP-5, both of which would be expected to have effects very different from localized and highly selective DA dysfunction. These notable differences underscore the risk of over-generalizing and highlight the significant impact that task requirements and experimental design can have on the results obtained.

Collectively, our present findings highlight the complex relationship between brain activity and brain function. While measurements of neural activity and neuromodulation offer valuable insights into how the brain responds to environmental stimuli or engages in cognition, caution is warranted when interpretating these neural signatures as a proxy for functional necessity (Tremblay et al. 2023). DA neuromodulation appears to have heterogeneous roles within basal ganglia circuits during learning and the expression of learning. Although phasic DA transients are evident across the striatum that respond dynamically to instrumental actions, causal manipulations underscore the indispensability of only some of these signals in shaping behavior. Importantly, this influence seems to be contingent upon the specific region and task being studied. Hence, we emphasize the need to complement correlational *in vivo* recordings with causal manipulations to determine the necessity of the derived signals in complex behavior (Robbins et al. 1990b; Szczypka et al. 2001b; Robinson et al. 2006; Yin et al. 2008b; Salamone and Correa 2012b; Darvas et al. 2014b; Stachniak et al. 2014; Fraser and Janak 2017b).

## Supplementary Figures

**Supplementary Fig. 1 | Dopamine (DA) signals in striatal regions at early (session 1) and late learning (session 19) aligned to four key task events (Trial Initiation, Correct and Incorrect Choice and Reward Collection).** (A) C57BL/6J mice underwent stereotaxic surgery for viral infusion and fiber optic implant in the nucleus accumbens (NAc), dorsomedial (DMS) and dorsolateral striatum (DLS). (B) Normalized mean DA signal in the NAc at early (session 1) and late learning (session 19) aligned to task events. Traces are plotted as 10s prior to and after the event (time 0 denoted by a dotted line). (C) Representative NAc DA dynamics (ΔF/F). After a 5 min habituation, subjects received a single injection of saline (i.p.) and DA was recorded for 1 hour. Cocaine (10mg.kg^-1^, i.p.) was then injected, followed by another hour. (D) Representative coronal brain section showing GFP immunoreactivity (GRAB_DA2M_, green; Hoechst, blue) within the NAc. (E) Normalized mean DA signal in the DMS at early (session 1) and late learning (session 19) aligned to task events. Traces are plotted as 10s prior to and after the event (time 0 denoted by a dotted line). (F) Representative DMS DA dynamics (ΔF/F). After a 5 min habituation, subjects received a single injection of saline (i.p.) and DA was recorded for 1 hour. Cocaine (10mg.kg^-1^, i.p.) was then injected, followed by another hour. (G) Representative coronal brain section showing GFP immunoreactivity (GRAB_DA2M_, green; Hoechst, blue) within the DMS. (H) Normalized mean DA signal in the DLS at early (session 1) and late learning (session 19) aligned to task events. Traces are plotted as 10s prior to and after the event (time 0 denoted by a dotted line). (I) Representative DLS DA dynamics (ΔF/F). After a 5 min habituation, subjects received a single injection of saline (i.p.) and DA was recorded for 1 hour. Cocaine (10mg.kg^-1^, i.p.) was then injected, followed by another hour. (J) Representative coronal brain section showing GFP immunoreactivity (GRAB_DA2M_, green; Hoechst, blue) within the DLS.

**Supplementary Fig. 2 | *In vivo* dopamine dynamics in the nucleus accumbens, dorsomedial and dorsolateral striatum differ from baseline and from one another most strongly in early relative to late learning.** (A) Graphical illustration of the quantifications in C-D, G-H, K-L, which compare the average max peak signals occurring following relevant task events relative to baseline signalling. (B) Graphical illustration of quantifications in E-F, I-J, M-O, which compare the average max peak signals between relevant task events. Early learning (Session 1) and late learning (Session 19) were compared for all measures. Relative to baseline, nucleus accumbens (NAc) dopamine (DA) was significantly different from baseline in early learning (C) but not late learning (D). Furthermore, task-relevant NAc-DA dynamics differed between themselves in early learning (E) but less so in late learning (F). Average DA max peak signals at correct choice was significantly different from baseline in the dorsomedial striatum (DMS), in both early (G) and late (H) learning. Furthermore, DMS-DA for correct choices were significantly different from incorrect choices and reward collection in early (I) but not late learning (J). Lastly, dorsolateral striatum (DLS) DA was significantly different from baseline in early learning (K), with significant differences at correct choice in late learning (L). Similarly, task-relevant DLS-DA dynamics differed between themselves in early learning (M), but only correct choice was significantly different from trial initiation in late learning (O). Data presented as Mean + SEM, 1-way RM-ANOVA X, * p<0.05, ** p<0.01, *** p<0.001, ****p<0.0001.

**Supplementary Fig. 3 | Ramp-like activity preceding choice is most prevalent in the nucleus accumbens, and not present in the dorsolateral striatum.** (A) Mean dopamine (DA) dynamics in the nucleus accumbens (NAc) prior to choice points in early learning (Session 1) and late learning (Session 19). Five second periods were compared, during a ‘baseline period’ and directly prior to choice, during a ‘ramp’ period. Area under curve (AUC) analyses between baseline and ramp periods in the NAc revealed a significant difference prior to making a correct choice in early (B) and late (C) learning, and prior to making an incorrect choice in early (D) and late (E) learning. (F) Mean DA dynamics in the dorsomedial striatum (DMS) prior to choice points in early and late learning were analyzed in the same way. AUC analyses between baseline and ramp periods in the DMS revealed a significant difference prior to making a correct choice in early (G) and late (H) learning, but no difference prior to making an incorrect choice at either timepoints (I-J). (K) Mean DA dynamics in the dorsolateral striatum (DLS) prior to choice points in early and late learning were similarly analyzed. AUC analyses between baseline and ramp periods in the DLS revealed no significant difference prior to making a correct or incorrect choice in either early or late learning (L-O). Data presented as Mean + SEM, Paired T-Test, * p<0.05, ** p<0.01.

**Supplementary Fig. 4 | Chemogenetic viral infection within the dorsolateral striatum enabled selective expression and inhibition of substantia nigra pars compacta dopamine neurons.** (A) Viral injection of a retrograde AAV expressing a Cre-dependent hM4D(Gi) (AAV-hSyn-DIO-hM4Di(Gi)-mCherry) into the dorsolateral striatum (DLS) induced selective colocalization of (B) TH and mCherry in the SN. (C) DREADD agonist clozapine N-Oxide (CNO) (1mg/kg, i.p.) reduced the percentage (%) of c-Fos+ positive cells in the DLS. (D) CNO inhibited dopamine neurons expressing mCherry in the SN. (E) Viral injection of a retrograde AAV expressing a Cre-dependent hM4D(Gi) (AAV-hSyn-DIO-hM4Di(Gi)-mCherry) into the DLS induced selective colocalization of TH and mCherry in the SNc. Representative coronal 20x and 40x magnification brain sections, Hoescht: blue; c-Fos: green; TH: purple; mCherry: red, scale bar 100 µm and 5 µm.

**Supplementary Fig. 5 | Chemogenetic viral infection within the nucleus accumbens enabled selective expression and inhibition of ventral tegmental area dopamine neurons.** (A) Viral injection of a retrograde AAV expressing a Cre-dependent hM4D(Gi) (AAV-hSyn-DIO-hM4Di(Gi)-mCherry) into the Nucleus Accumbens (NAc) induced selective colocalization of TH and mCherry (B) in the ventral tegmental area (VTA). C) DREADD agonist clozapine N-Oxide (CNO) (1mg/kg, i.p.) reduced the percentage (%) of c-Fos+ positive cells in the NAc. (D) CNO inhibited dopamine neurons expressing mCherry in the VTA. (E) Viral injection of a retrograde AAV expressing a Cre-dependent hM4D(Gi) (AAV-hSyn-DIO-hM4Di(Gi)-mCherry) into the NAc induced selective colocalization of TH and mCherry in the VTA. Representative coronal 20x and 40x magnification brain sections, Hoescht: blue; c-Fos: green; TH: purple; mCherry: red, scale bar 100 µm and 5 µm.

**Supplementary Fig. 6 | Inhibition of DLS-projecting nigrostriatal and NAc-projecting mesolimbic dopamine has minimal impact on expression of previous learning.** (A) DAT-IRES-Cre mice underwent stereotaxic surgery for viral infusion of a retrograde inhibitory hM4D(Gi) DREADD or the mCherry control transgene into either the dorsolateral striatum (which retroactively targeted the nigrostriatal dopamine pathway) or the nucleus accumbens (which retroactively targeted the mesolimbic dopamine pathway). Following post-operative recovery, subjects ran on the Visuomotor Conditional Learning (VMCL) training task until they attained high accuracy, and then received acute systemic injections of either saline control or CNO (1mg/kg, i.p.) 30 minutes prior to task onset. Data are presented as the average across two sessions, order counterbalanced. No significant difference was observed in the number of sessions to reach a criterion of >77% on the VMCL Training task for 2 consecutive sessions in pre-manipulation comparisons between experimental groups and their respected controls (B-C). Once criterion was met, supplementary measures of VMCL performance was compared on saline versus CNO, including perseveration index (D-E), number of correction trials (F-G), % missed trials (H-I), and latencies to collect reward (J-K) and make a correct (L-M) or incorrect choices (N-O). To our surprise, the only significant difference revealed was a mild increase in the % missed trials across both drug conditions for the mesolimbic DA inhibition group relative to their controls (I). This increase was not present in pre-manipulation testing (data not shown). Data presented as Mean + SEM, 2-way RM-ANOVA, * p<0.05.

**Supplementary Fig. 7 | Inhibition of DLS-projecting dopamine but not NAc-projecting mesolimbic dopamine led to selective impairment in visuomotor conditional learning.** (A) DAT-IRES-Cre mice underwent stereotaxic surgery for viral infusion of a retrograde inhibitory DREADD hM4D(Gi) or a mCherry control transgene into either the dorsolateral striatum (which retroactively targeted the nigrostriatal dopamine pathway) or the nucleus accumbens (which retroactively targeted the mesolimbic pathway). During Visuomotor Conditional Learning (VMCL) Test, subjects received 20 daily injections of clozapine N-Oxide (CNO) (1mg/kg, i.p.) 30 minutes prior to task onset. Relative to their controls, the DLS-projecting nigrostriatal DA inhibition and NAc-projecting mesolimbic DA inhibition groups were compared on secondary measures of the VMCL task, including perseveration index (B-C), number of correction trials (D-E), % missed trials (F-G, and latencies to collect reward (H-I), and make correct (J-K) and incorrect choices (L-M). Relative to their controls, this revealed an elevated perseveration index (B) and longer incorrect touch latency (L) in the DLS-projecting nigrostriatal DA inhibition group, but no impairments in the NAc-projecting mesolimbic DA inhibition group. Furthermore, unblocked sessions of % correct responses revealed a significant impairment in task acquisition in the DLS-projecting nigrostriatal DA inhibition group and a failure to display asymptotic behaviour in the last 6 sessions (defined as no difference between consecutive mean percent correct scores). No difference was found between the NAc-projecting mesolimbic DA inhibition group (O) their controls, with both groups displaying asymptotic behaviour in the last 6 sessions. Data presented as Mean + SEM, 2-way RM-ANOVA, * p<0.05 ** p<0.01.

**Supplementary Fig. 8 | DLS-projecting nigrostriatal, but not NAc-projecting mesolimbic dopamine inhibition has a long-lasting impact on visuomotor conditional learning test performance.** (A) DAT-IRES-Cre mice underwent stereotaxic surgery for viral infusion of the retrograde inhibitory DREADD hM4D(Gi) or the mCherry control into either the dorsolateral striatum (which retroactively targeted the nigrostriatal dopamine pathway) or the nucleus accumbens (which retroactively targeted the mesolimbic pathway). Following the acquisition of Visuomotor Conditional Learning (VMCL) Test, where subjects received 20 daily injections of clozapine N-Oxide (CNO) (1mg/kg, i.p.), subjects were then introduced to 4 sessions of inhibition relief (saline), followed by 4 sessions of reinstatement (CNO) to assess the long-lasting effects of dopaminergic blockade. Data is presented as the average of 4 sessions, comparing block 5 of VMCL test, saline inhibition relief and CNO reinstatement. DLS-projecting nigrostriatal and NAc-projecting mesolimbic DA inhibition groups were compared to their relative controls and assessed secondary measures of the VMCL test: perseveration index (B-C), number of correction trials (D-E), % missed trials (F-G, and latencies to collect reward (H-I) and make correct (J-K) and incorrect choices (L-M). This revealed significant and long-lasting impairments of DLS-projecting nigrostriatal DA inhibition, but no effect following NAc-projecting mesolimbic DA inhibition. Data presented as Mean + SEM, 2-way RM-ANOVA, * p<0.05, ** p<0.01.

## Acknowledgements

We thank Jue Fan, Chris Fodor and the Animal Care and Veterinary Services (ACVS) staff at Western University for animal care. OPL received a Natural Science and Engineering Research Council of Canada (NSERC) graduate scholarship along with a Canadian Institutes for Health Research (CIHR) graduate scholarship. MS received funding support from the Canada First Research Excellence Fund Accelerator Awards (BrainsCAN). M.A.M.P., V.F.P., L.M.S., T.J.B., received support from CIHR (PJT 426966, 162432, 159781, 169101), NSERC (402524-2013 RGPIN; 03592-2021 RGPIN; 06711-2019 RGPIN), Canada First Research Excellence Fund Accelerator Awards (BrainsCAN), TRanslational Initiative to DE-risk NeuroTherapeutics (TRIDENT) New Frontiers in Research Fund (NFRF), Brain Canada, Canada Foundation for Innovation (CFI) fund (43110, 36569, 37479), and Ontario Research Fund. M.A.M.P is a Tier I Canada Research Chair in Neurochemistry of Dementia. L.M.S. is a Tier I Canada Research Chair in Translational Cognitive Neuroscience. T.J.B is a Western Research Chair.

## Competing interests

T.J.B and L.M.S have established a series of targeted cognitive tests for animals, administered via touchscreen with a custom environment known as the “Bussey-Saksida touchscreen chamber”. Cambridge Enterprise, the technology transfer office of the University of Cambridge, supported commercialisation of the Bussey-Saksida chamber, culminating in a license to Campden Instruments. Any financial compensation received from commercialisation of the technology is fully invested in further touchscreen development and/or maintenance. All other authors declare no competing interests.

## Author Contributions Section

O.P.L, M.S., L.M.S. and T.J.B designed experiments. O.P.L, M.M.F.M., M.S., H.R., S.P., A.C., C.A.L. and N.D. performed experiments. O.P.L analyzed the data. D.P. designed the python-based code to extract and analyse fiber photometry data and both O.P.L and D.P. designed the behavioural schedules on Abet software. O.P.L, M.M.F.M, L.M.S. and T.J.B wrote the manuscript.

## Notes

### Summary of Updates

Minor revisions have been made throughout the manuscript.

## References

Armbruster BN, Li X, Pausch MH, et al (2007) Evolving the lock to fit the key to create a family of G protein-coupled receptors potently activated by an inert ligand. Proc Natl Acad Sci U S A 104:5163–5168. 10.1073/PNAS.0700293104/SUPPL_FILE/00293FIG8.PDF

Ashby FG, Ennis JM, Spiering BJ (2007) A neurobiological theory of automaticity in perceptual categorization. Psychol Rev 114:632–656. 10.1037/0033-295X.114.3.632

Atallah HE, Lopez-Paniagua D, Rudy JW, O’Reilly RC (2007) Separate neural substrates for skill learning and performance in the ventral and dorsal striatum. Nat Neurosci 10:126–131. 10.1038/NN1817

Bayer HM, Glimcher PW (2005) Midbrain dopamine neurons encode a quantitative reward prediction error signal. Neuron 47:129–141. 10.1016/j.neuron.2005.05.020

Beninger RJ, Hahn BL (1983) Pimozide blocks establishment but not expression of amphetamine-produced environment-specific conditioning. Science (1979) 220:1304–1306. 10.1126/SCIENCE.6857251

Berger TW, Thompson RF (1978) Neuronal plasticity in the limbic system during classical conditioning of the rabbit nictitating membrane response. I. The hippocampus. Brain Res 145:323–346. 10.1016/0006-8993(78)90866-1

Bermudez MA, Schultz W (2014) Timing in reward and decision processes. Philosophical Transactions of the Royal Society B: Biological Sciences 369:20120468. 10.1098/RSTB.2012.0468

Bernheimer H, Birkmayer W, Hornykiewicz O, et al (1973) Brain dopamine and the syndromes of Parkinson and Huntington Clinical, morphological and neurochemical correlations. J Neurol Sci 20:415–455. 10.1016/0022-510X(73)90175-5

Brain M, Krupa DJ, Thompson JK, Thompson RF (1993) Localization of a Memory Trace in the Mammalian Brain. Source: Science 260:989–991

Brown HD, Mccutcheon JE, Cone JJ, et al (2011) Primary food reward and reward-predictive stimuli evoke different patterns of phasic dopamine signaling throughout the striatum. Eur J Neurosci 34:1997–2006. 10.1111/J.1460-9568.2011.07914.X

Cachope R, Mateo Y, Mathur BN, et al (2012) Selective activation of cholinergic interneurons enhances accumbal phasic dopamine release: setting the tone for reward processing. Cell Rep 2:33–41. 10.1016/J.CELREP.2012.05.011

Calipari ES, Huggins KN, Mathews TA, Jones SR (2012) Conserved dorsal-ventral gradient of dopamine release and uptake rate in mice, rats and rhesus macaques. Neurochem Int 61:986–991. 10.1016/J.NEUINT.2012.07.008

Carelli RM, Wolske M, West MO (1997) Loss of Lever Press-Related Firing of Rat Striatal Forelimb Neurons after Repeated Sessions in a Lever Pressing Task. The Journal of Neuroscience 17:1804. 10.1523/JNEUROSCI.17-05-01804.1997

Centonze D, Grande C, Saulle E, et al (2003) Distinct Roles of D1 and D5 Dopamine Receptors in Motor Activity and Striatal Synaptic Plasticity. The Journal of Neuroscience 23:8506. 10.1523/JNEUROSCI.23-24-08506.2003

Centonze D, Gubellini P, Picconi B, et al (1999) Unilateral dopamine denervation blocks corticostriatal LTP. J Neurophysiol 82:3575–3579. 10.1152/JN.1999.82.6.3575/ASSET/IMAGES/LARGE/9K1190587002.JPEG

Chen APF, Malgady JM, Chen L, et al (2022) Nigrostriatal dopamine pathway regulates auditory discrimination behavior. Nat Commun 13:. 10.1038/S41467-022-33747-2

Choi WY, Balsam PD, Horvitz JC (2005) Extended habit training reduces dopamine mediation of appetitive response expression. J Neurosci 25:6729–6733. 10.1523/JNEUROSCI.1498-05.2005

Choi WY, Morvan C, Balsam PD, Horvitz JC (2009) Dopamine D1 and D2 Antagonist Effects on Response Likelihood and Duration. Behavioral Neuroscience 123:1279–1287. 10.1037/A0017702

Collins AL, Greenfield VY, Bye JK, et al (2016a) Dynamic mesolimbic dopamine signaling during action sequence learning and expectation violation. Sci Rep 6:20231. 10.1038/srep20231

Collins AL, Greenfield VY, Bye JK, et al (2016b) Dynamic mesolimbic dopamine signaling during action sequence learning and expectation violation. Scientific Reports 2016 6:1 6:1–15. 10.1038/srep20231

Collins AL, Saunders BT (2020) Heterogeneity in striatal dopamine circuits: Form and function in dynamic reward seeking HHS Public Access. J Neurosci Res 98:1046–1069. 10.1002/jnr.24587

Costa RM, Lin SC, Sotnikova TDD, et al (2006) Rapid alterations in corticostriatal ensemble coordination during acute dopamine-dependent motor dysfunction. Neuron 52:359–369. 10.1016/J.NEURON.2006.07.030

Cox J, Witten IB (2019) Striatal circuits for reward learning and decision-making. Nature Reviews Neuroscience 2019 20:8 20:482–494. 10.1038/s41583-019-0189-2

Cragg SJ, Hille CJ, Greenfield SA (2000) Dopamine Release and Uptake Dynamics within Nonhuman Primate Striatum In Vitro. The Journal of Neuroscience 20:8209. 10.1523/JNEUROSCI.20-21-08209.2000

Darvas M, Palmiter RD (2009) Restriction of dopamine signaling to the dorsolateral striatum is sufficient for many cognitive behaviors. Proc Natl Acad Sci U S A 106:14664–14669. 10.1073/PNAS.0907299106;WEBSITE:WEBSITE:PNAS-SITE;ISSUE:ISSUE:DOI

Darvas M, Wunsch AM, Gibbs JT, Palmiter RD (2014a) Dopamine dependency for acquisition and performance of Pavlovian conditioned response. Proc Natl Acad Sci U S A 111:2764–2769. 10.1073/PNAS.1400332111/SUPPL_FILE/PNAS.201400332SI.PDF

Darvas M, Wunsch AM, Gibbs JT, Palmiter RD (2014b) Dopamine dependency for acquisition and performance of Pavlovian conditioned response. Proc Natl Acad Sci U S A 111:2764–2769. 10.1073/pnas.1400332111

Dauer W, Przedborski S (2003) Parkinson’s Disease: Mechanisms and Models. Neuron 39:889–909. 10.1016/S0896-6273(03)00568-3

Delotterie D, Mathis C, Cassel JC, et al (2014) Optimization of Touchscreen-Based Behavioral Paradigms in Mice: Implications for Building a Battery of Tasks Taxing Learning and Memory Functions. PLoS One 9:e100817. 10.1371/JOURNAL.PONE.0100817

Delotterie DF, Mathis C, Cassel JC, et al (2015) Touchscreen tasks in mice to demonstrate differences between hippocampal and striatal functions. Neurobiol Learn Mem 120:16–27. 10.1016/J.NLM.2015.02.007

Dextera DT, Jenner P (2013) Parkinson disease: from pathology to molecular disease mechanisms. Free Radic Biol Med 62:132–144. 10.1016/J.FREERADBIOMED.2013.01.018

Dickinson A (1985) Actions and habits: the development of behavioural autonomy. Philosophical Transactions of the Royal Society of London B, Biological Sciences 308:67–78. 10.1098/RSTB.1985.0010

Engelhard B, Finkelstein J, Cox J, et al (2019) Specialized coding of sensory, motor and cognitive variables in VTA dopamine neurons. Nature 2019 570:7762 570:509–513. 10.1038/s41586-019-1261-9

Eshel N, Touponse GC, Wang AR, et al (2023) Striatal dopamine integrates cost, benefit, and motivation. Neuron. 10.1016/J.NEURON.2023.10.038

Exley R, Cragg SJ (2008) Presynaptic nicotinic receptors: a dynamic and diverse cholinergic filter of striatal dopamine neurotransmission. Br J Pharmacol 153:S283–S297. 10.1038/SJ.BJP.0707510

Farrell K, Lak A, Saleem AB (2022) Midbrain dopamine neurons signal phasic and ramping reward prediction error during goal-directed navigation. Cell Rep 41:. 10.1016/J.CELREP.2022.111470

Fraser KM, Janak PH (2017a) Long-lasting contribution of dopamine in the nucleus accumbens core, but not dorsal lateral striatum, to sign-tracking. Eur J Neurosci 46:2047. 10.1111/EJN.13642

Fraser KM, Janak PH (2017b) Long-lasting contribution of dopamine in the nucleus accumbens core, but not dorsal lateral striatum, to sign-tracking. European Journal of Neuroscience 46:2047–2055. 10.1111/ejn.13642

Galea LAM, Choleris E, Albert AYK, et al (2020) The promises and pitfalls of sex difference research. Front Neuroendocrinol 56:. 10.1016/J.YFRNE.2019.100817

Garrett MC, Soares-da-Silva P (1992) Increased cerebrospinal fluid dopamine and 3,4-dihydroxyphenylacetic acid levels in Huntington’s disease: evidence for an overactive dopaminergic brain transmission. J Neurochem 58:101–106. 10.1111/J.1471-4159.1992.TB09283.X

Garris PA, Ciolkowski EL, Wightman RM (1994a) Heterogeneity of evoked dopamine overflow wihin the striatal and striatoamygdaloid regions. Neuroscience 59:417–427. 10.1016/0306-4522(94)90606-8

Garris PA, Ciolkowski EL, Wightman RM (1994b) Heterogeneity of evoked dopamine overflow within the striatal and striatoamygdaloid regions. Neuroscience 59:417–427. 10.1016/0306-4522(94)90606-8

Gomez JL, Bonaventura J, Lesniak W, et al (2017) Chemogenetics revealed: DREADD occupancy and activation via converted clozapine. Science 357:503–507. 10.1126/SCIENCE.AAN2475

Gowrishankar R, Gresch PJ, Davis GL, et al (2018) Region-Specific Regulation of Presynaptic Dopamine Homeostasis by D2 Autoreceptors Shapes the In Vivo Impact of the Neuropsychiatric Disease-Associated DAT Variant Val559. Journal of Neuroscience 38:5302–5312. 10.1523/JNEUROSCI.0055-18.2018

Groenewegen HJ, Uylings HBM (2010) Organization of Prefrontal-Striatal Connections. Handb Behav Neurosci 20:353–365. 10.1016/B978-0-12-374767-9.00020-2

Hadj-Bouziane F, Boussaoud D (2003) Neuronal activity in the monkey striatum during conditional visuomotor learning. Exp Brain Res 153:190–196. 10.1007/S00221-003-1592-4

Hallett PJ, Spoelgen R, Hyman BT, et al (2006) Dopamine D1 Activation Potentiates Striatal NMDA Receptors by Tyrosine Phosphorylation-Dependent Subunit Trafficking. Journal of Neuroscience 26:4690–4700. 10.1523/JNEUROSCI.0792-06.2006

Hamid AA, Frank MJ, Moore CI (2021) Wave-like dopamine dynamics as a mechanism for spatiotemporal credit assignment. Cell 184:2733–2749.e16. 10.1016/J.CELL.2021.03.046

Hamid AA, Pettibone JR, Mabrouk OS, et al (2015) Mesolimbic dopamine signals the value of work. Nature Neuroscience 2016 19:1 19:117–126. 10.1038/nn.4173

Hart AS, Rutledge RB, Glimcher PW, Phillips PEM (2014) Phasic Dopamine Release in the Rat Nucleus Accumbens Symmetrically Encodes a Reward Prediction Error Term. Journal of Neuroscience 34:698–704. 10.1523/JNEUROSCI.2489-13.2014

Hayes HA, Hunsaker N, Dibble LE (2015) Implicit Motor Sequence Learning in Individuals with Parkinson Disease: A Meta-Analysis. J Parkinsons Dis 5:549–560. 10.3233/JPD-140441

Heymann G, Jo YS, Reichard KL, et al (2020) Synergy of Distinct Dopamine Projection Populations in Behavioral Reinforcement. Neuron 105:909–920.e5. 10.1016/J.NEURON.2019.11.024

Hiebert NM, Owen AM, Ganjavi H, et al (2019) Dorsal striatum does not mediate feedback-based, stimulus-response learning: An event-related fMRI study in patients with Parkinson’s disease tested on and off dopaminergic therapy. Neuroimage 185:455–470. 10.1016/J.NEUROIMAGE.2018.10.045

Hiebert NM, Owen AM, Seergobin KN, MacDonald PA (2017) Dorsal striatum mediates deliberate decision making, not late-stage, stimulus-response learning. Hum Brain Mapp 38:6133–6156. 10.1002/HBM.23817

Hirsch E, Graybiel AM, Agid YA (1988) Melanized dopaminergic neurons are differentially susceptible to degeneration in Parkinson’s disease. Nature 334:345–348. 10.1038/334345A0

Holland PC (2008) Cognitive versus stimulus-response theories of learning. Learn Behav 36:227. 10.3758/LB.36.3.227

Horner AE, Heath CJ, Hvoslef-Eide M, et al (2013) The touchscreen operant platform for testing learning and memory in rats and mice. Nature Protocols 2013 8:10 8:1961–1984. 10.1038/nprot.2013.122

Horvitz JC (2009) Stimulus–response and response–outcome learning mechanisms in the striatum. Behavioural Brain Research 199:129–140. 10.1016/J.BBR.2008.12.014

Horvitz JC (2002) Dopamine gating of glutamatergic sensorimotor and incentive motivational input signals to the striatum. Behavioural Brain Research 137:65–74. 10.1016/S0166-4328(02)00285-1

Horvitz JC, Ettenberg A (1991) Conditioned Incentive Properties of a Food-Paired Conditioned Stimulus Remain Intact During Dopamine Receptor Blockade. Behavioral Neuroscience 105:536–541. 10.1037/0735-7044.105.4.536

Houk JC, Wise SP (1995) Distributed modular architectures linking basal ganglia, cerebellum, and cerebral cortex: their role in planning and controlling action. Cereb Cortex 5:95–110. 10.1093/CERCOR/5.2.95

Howe MW, Dombeck DA (2016) Rapid signaling in distinct dopaminergic axons during locomotion and reward. Nature 535:505. 10.1038/NATURE18942

Howe MW, Tierney PL, Sandberg SG, et al (2013) Prolonged dopamine signalling in striatum signals proximity and value of distant rewards. Nature 500:575–579. 10.1038/NATURE12475

Joel D, Niv Y, Ruppin E (2002) Actor–critic models of the basal ganglia: new anatomical and computational perspectives. Neural Networks 15:535–547. 10.1016/S0893-6080(02)00047-3

Jones IW, Paul Bolam J, Wonnacott S (2001) Presynaptic localisation of the nicotinic acetylcholine receptor beta2 subunit immunoreactivity in rat nigrostriatal dopaminergic neurones. J Comp Neurol 439:235–247. 10.1002/CNE.1345

Jones SR, Garris PA, Kilts CD, Wightman RM (1995) Comparison of dopamine uptake in the basolateral amygdaloid nucleus, caudate-putamen, and nucleus accumbens of the rat. J Neurochem 64:2581–2589. 10.1046/J.1471-4159.1995.64062581.X

Kalia L V., Lang AE (2015) Parkinson’s disease. Lancet 386:896–912. 10.1016/S0140-6736(14)61393-3

Keiflin R, Janak PH (2015) Dopamine prediction errors in reward learning and addiction: from theory to neural circuitry. Neuron 88:247. 10.1016/J.NEURON.2015.08.037

Kerr JND, Wickens JR (2001) Dopamine D-1/D-5 receptor activation is required for long-term potentiation in the rat neostriatum in vitro. J Neurophysiol 85:117–124. 10.1152/JN.2001.85.1.117

Kim HGR, Malik AN, Mikhael JG, et al (2020) A Unified Framework for Dopamine Signals across Timescales. Cell 183:1600–1616.e25. 10.1016/J.CELL.2020.11.013

Kljakic O, Janíčková H, Skirzewski M, et al (2022a) Functional dissociation of behavioral effects from acetylcholine and glutamate released from cholinergic striatal interneurons. The FASEB Journal 36:e22135. 10.1096/FJ.202101425R

Kljakic O, Janíčková H, Skirzewski M, et al (2022b) Functional dissociation of behavioral effects from acetylcholine and glutamate released from cholinergic striatal interneurons. The FASEB Journal 36:. 10.1096/FJ.202101425R

Kljakic O, Janíčková H, Skirzewski M, et al (2022c) Functional dissociation of behavioral effects from acetylcholine and glutamate released from cholinergic striatal interneurons. The FASEB Journal 36:. 10.1096/FJ.202101425R

Knopman D, Nissen MJ (1991) Procedural learning is impaired in Huntington’s disease: Evidence from the serial reaction time task. Neuropsychologia 29:245–254. 10.1016/0028-3932(91)90085-M

Krashes MJ, Koda S, Ye CP, et al (2011) Rapid, reversible activation of AgRP neurons drives feeding behavior in mice. J Clin Invest 121:1424. 10.1172/JCI46229

Krishnan S, Heer C, Cherian C, Sheffield MEJ (2022) Reward expectation extinction restructures and degrades CA1 spatial maps through loss of a dopaminergic reward proximity signal. Nature Communications 2022 13:1 13:1–19. 10.1038/s41467-022-34465-5

Lakens D (2013) Calculating and reporting effect sizes to facilitate cumulative science: A practical primer for t-tests and ANOVAs. Front Psychol 4:62627. 10.3389/FPSYG.2013.00863/ABSTRACT

Lawrence AD, Sahakian BJ, Rogers RD, et al (1999) Discrimination, reversal, and shift learning in Huntington’s disease: mechanisms of impaired response selection. Neuropsychologia 37:1359–1374. 10.1016/S0028-3932(99)00035-4

Legaria AA, Matikainen-Ankney BA, Yang B, et al (2022) Fiber photometry in striatum reflects primarily nonsomatic changes in calcium. Nature Neuroscience 2022 25:9 25:1124–1128. 10.1038/s41593-022-01152-z

MacLaren DAA, Browne RW, Shaw JK, et al (2016) Clozapine N-Oxide Administration Produces Behavioral Effects in Long-Evans Rats: Implications for Designing DREADD Experiments. eNeuro 3:. 10.1523/ENEURO.0219-16.2016

Malvaez M, Wassum KM (2018) Regulation of habit formation in the dorsal striatum. Curr Opin Behav Sci 20:67–74. 10.1016/J.COBEHA.2017.11.005

Mar AC, Horner AE, Nilsson SRO, et al (2013) The touchscreen operant platform for assessing executive function in rats and mice. Nat Protoc 8:1985–2005. 10.1038/NPROT.2013.123

Markowitz JE, Gillis WF, Beron CC, et al (2018) The Striatum Organizes 3D Behavior via Moment-to-Moment Action Selection. Cell 174:44–58.e17. 10.1016/J.CELL.2018.04.019

Mechawar N Cholinergic innervation in adult rat cerebral cortex: A quantitative immunocytochemical description. J Comp Neurol 428:

Menegas W, Babayan BM, Uchida N, Watabe-Uchida M (2017) Opposite initialization to novel cues in dopamine signaling in ventral and posterior striatum in mice. Elife 6:. 10.7554/ELIFE.21886

Mi TM, Zhang W, McKeown MJ, Chan P (2021) Impaired Formation and Expression of Goal-Directed and Habitual Control in Parkinson’s Disease. Front Aging Neurosci 13:734807. 10.3389/FNAGI.2021.734807

Miller LR, Marks C, Becker JB, et al (2017) Considering sex as a biological variable in preclinical research. The FASEB Journal 31:29. 10.1096/FJ.201600781R

Mohebi A, Pettibone JR, Hamid AA, et al (2019) Dissociable dopamine dynamics for learning and motivation. Nature 2019 570:7759 570:65–70. 10.1038/s41586-019-1235-y

Mohebi A, Wei W, Pelattini L, et al (2024a) Dopamine transients follow a striatal gradient of reward time horizons. Nature Neuroscience 2024 27:4 27:737–746. 10.1038/s41593-023-01566-3

Mohebi A, Wei W, Pelattini L, et al (2024b) Dopamine transients follow a striatal gradient of reward time horizons. Nature Neuroscience 2024 1–10. 10.1038/s41593-023-01566-3

Musall S, Kaufman MT, Juavinett AL, et al (2019) Single-trial neural dynamics are dominated by richly varied movements. Nature Neuroscience 2019 22:10 22:1677–1686. 10.1038/s41593-019-0502-4

Nolan SO, Zachry JE, Johnson AR, et al (2020) Direct dopamine terminal regulation by local striatal microcircuitry. J Neurochem 155:475. 10.1111/JNC.15034

O’Donnell P (2003) Dopamine gating of forebrain neural ensembles. European Journal of Neuroscience 17:429–435. 10.1046/J.1460-9568.2003.02463.X

Olds J, Milner P (1954) Positive reinforcement produced by electrical stimulation of septal area and other regions of rat brain. J Comp Physiol Psychol 47:419–427. 10.1037/H0058775

Olson M, Lockhart TE, Lieberman A (2019) Motor Learning Deficits in Parkinson’s Disease (PD) and Their Effect on Training Response in Gait and Balance: A Narrative Review. Front Neurol 10:. 10.3389/FNEUR.2019.00062

O’Reilly RC, Frank MJ (2006) Making working memory work: a computational model of learning in the prefrontal cortex and basal ganglia. Neural Comput 18:283–328. 10.1162/089976606775093909

Ostlund SB, Balleine BW (2008) On habits and addiction: An associative analysis of compulsive drug seeking. Drug Discov Today Dis Models 5:235. 10.1016/J.DDMOD.2009.07.004

Piantadosi PT, Princz-Lebel O, Skirzewski M, et al (2025) Integrating optical neuroscience tools into touchscreen operant systems. Nature Protocols 2025 1–35. 10.1038/s41596-025-01143-x

Poulin JF, Caronia G, Hofer C, et al (2018) Mapping projections of molecularly defined dopamine neuron subtypes using intersectional genetic approaches. Nature Neuroscience 2018 21:9 21:1260–1271. 10.1038/s41593-018-0203-4

Reading PJ, Dunnett SB, Robbins TW (1991) Dissociable roles of the ventral, medial and lateral striatum on the acquisition and performance of a complex visual stimulus-response habit. Behavioural Brain Research 45:147–161. 10.1016/S0166-4328(05)80080-4

Reynolds JNJ, Hyland BI, Wickens JR (2001) A cellular mechanism of reward-related learning. Nature 2001 413:6851 413:67–70. 10.1038/35092560

Robbins T, Giardini V, Jones G, Sahakian B (1990a) Effects of dopamine depletion from the caudate-putamen and nucleus accumbens septi on the acquisition and performance of a conditional discrimination task. Behavioural Brain Research 38:243–261

Robbins TW, Giardini V, Jones GH, et al (1990b) Effects of dopamine depletion from the caudate-putamen and nucleus accumbens septi on the acquisition and performance of a conditional discrimination task. Behavioural brain research 38:243–261. 10.1016/0166-4328(90)90179-I

Robinson S, Rainwater AJ, Hnasko TS, Palmiter RD (2007) Viral restoration of dopamine signaling to the dorsal striatum restores instrumental conditioning to dopamine-deficient mice. Psychopharmacology (Berl) 191:567–578. 10.1007/S00213-006-0579-9

Robinson S, Rainwater AJ, Hnasko TS, Palmiter RD (2006) Viral restoration of dopamine signaling to the dorsal striatum restores instrumental conditioning to dopamine-deficient mice. Psychopharmacology 2006 191:3 191:567–578. 10.1007/s00213-006-0579-9

Rothblat DS, Schneider JS (1997) Regionally specific effects of haloperidol and clozapine on dopamine reuptake in the striatum. Neurosci Lett 228:119–122. 10.1016/S0304-3940(97)00377-7

Runegaard AH, Fitzpatrick CM, Woldbye DPD, et al (2019) Modulating Dopamine Signaling and Behavior with Chemogenetics: Concepts, Progress, and Challenges. Pharmacol Rev 71:123–156. 10.1124/PR.117.013995

Runegaard AH, Sørensen AT, Fitzpatrick CM, et al (2018) Locomotor- and Reward-Enhancing Effects of Cocaine Are Differentially Regulated by Chemogenetic Stimulation of Gi-Signaling in Dopaminergic Neurons. eNeuro 5:. 10.1523/ENEURO.0345-17.2018

Salamone JD, Correa M (2012a) The Mysterious Motivational Functions of Mesolimbic Dopamine. Neuron 76:470–485. 10.1016/J.NEURON.2012.10.021

Salamone JD, Correa M (2012b) The Mysterious Motivational Functions of Mesolimbic Dopamine. Neuron 76:470–485. 10.1016/j.neuron.2012.10.021

Salinas AG, Lee JO, Augustin SM, et al (2023a) Distinct sub-second dopamine signaling in dorsolateral striatum measured by a genetically-encoded fluorescent sensor. Nature Communications 2023 14:1 14:1–16. 10.1038/s41467-023-41581-3

Salinas AG, Lee JO, Augustin SM, et al (2023b) Distinct sub-second dopamine signaling in dorsolateral striatum measured by a genetically-encoded fluorescent sensor. Nat Commun 14:5915. 10.1038/s41467-023-41581-3

Schmaltz LW, Theios J (1972) Acquisition and extinction of a classically conditioned response in hippocampectomized rabbits (Oryctolagus cuniculus). J Comp Physiol Psychol 79:328–333. 10.1037/H0032531

Schultz W (2016) Dopamine reward prediction error coding. Dialogues Clin Neurosci 18:23. 10.31887/DCNS.2016.18.1/WSCHULTZ

Schultz W (2013) Updating dopamine reward signals. Curr Opin Neurobiol 23:229–238. 10.1016/J.CONB.2012.11.012

Schultz W (2010) Dopamine signals for reward value and risk: Basic and recent data. Behavioral and Brain Functions 6:1–9. 10.1186/1744-9081-6-24/FIGURES/2

Schultz W, Dayan P, Montague PR (1997) A neural substrate of prediction and reward. Science (1979) 275:1593–1599. 10.1126/SCIENCE.275.5306.1593;JOURNAL:JOURNAL:SCIENCE;WGROUP:STRING:PUBLICATION

Seiler JL, Cosme C V., Sherathiya VN, et al (2022a) Dopamine signaling in the dorsomedial striatum promotes compulsive behavior. Current Biology 32:1175–1188.e5. 10.1016/j.cub.2022.01.055

Seiler JL, Cosme C V., Sherathiya VN, et al (2022b) Dopamine signaling in the dorsomedial striatum promotes compulsive behavior. Curr Biol 32:1175–1188.e5. 10.1016/J.CUB.2022.01.055

Shin JH, Adrover MF, Wess J, Alvarez VA (2015) Muscarinic regulation of dopamine and glutamate transmission in the nucleus accumbens. Proc Natl Acad Sci U S A 112:8124–8129. 10.1073/PNAS.1508846112/SUPPL_FILE/PNAS.201508846SI.PDF

Skirzewski M, Princz-Lebel O, German-Castelan L, et al (2022) Continuous cholinergic-dopaminergic updating in the nucleus accumbens underlies approaches to reward-predicting cues. Nature Communications 2022 13:1 13:1–21. 10.1038/s41467-022-35601-x

Solomon PR, Moore JW (1975) Latent inhibition and stimulus generalization of the classically conditioned nictitating membrane response in rabbits (Oryctolagus cuniculus) following dorsal hippocampal ablation. J Comp Physiol Psychol 89:1192–1203. 10.1037/H0077183

Sotak BN, Hnasko TS, Robinson S, et al (2005) Dysregulation of dopamine signaling in the dorsal striatum inhibits feeding. Brain Res 1061:88–96. 10.1016/J.BRAINRES.2005.08.053

Spokes EGS (1980) Neurochemical alterations in Huntington’s chorea: a study of post-mortem brain tissue. Brain 103:179–210. 10.1093/BRAIN/103.1.179

Stachniak TJ, Ghosh A, Sternson SM (2014) Chemogenetic Synaptic Silencing of Neural Circuits Localizes a Hypothalamus→Midbrain Pathway for Feeding Behavior. Neuron 82:797–808. 10.1016/j.neuron.2014.04.008

Sulzer D, Cragg SJ, Rice ME (2016) Striatal dopamine neurotransmission: regulation of release and uptake. Basal Ganglia 6:123–148. 10.1016/J.BAGA.2016.02.001

Sun F, Zhou J, Dai B, et al (2020) Next-generation GRAB sensors for monitoring dopaminergic activity in vivo. Nat Methods 17:1156. 10.1038/S41592-020-00981-9

Szczypka MS, Kwok K, Brot MD, et al (2001a) Dopamine production in the caudate putamen restores feeding in dopamine-deficient mice. Neuron 30:819–828. 10.1016/S0896-6273(01)00319-1

Szczypka MS, Kwok K, Brot MD, et al (2001b) Dopamine production in the caudate putamen restores feeding in dopamine-deficient mice. Neuron 30:819–828. 10.1016/S0896-6273(01)00319-1

Threlfell S, Clements MA, Khodai T, et al (2010) Striatal muscarinic receptors promote activity dependence of dopamine transmission via distinct receptor subtypes on cholinergic interneurons in ventral versus dorsal striatum. J Neurosci 30:3398–3408. 10.1523/JNEUROSCI.5620-09.2010

Threlfell S, Lalic T, Platt NJ, et al (2012a) Striatal Dopamine Release Is Triggered by Synchronized Activity in Cholinergic Interneurons. Neuron 75:58–64. 10.1016/J.NEURON.2012.04.038

Threlfell S, Lalic T, Platt NJ, et al (2012b) Striatal dopamine release is triggered by synchronized activity in cholinergic interneurons. Neuron 75:58–64. 10.1016/J.NEURON.2012.04.038

Threlfell S, Lalic T, Platt NJ, et al (2012c) Striatal dopamine release is triggered by synchronized activity in cholinergic interneurons. Neuron 75:58–64. 10.1016/J.NEURON.2012.04.038

Tremblay S, Testard C, Inchauspé J, Petrides M Epiphenomenal neural activity in the primate cortex. 10.1101/2022.09.12.506984

Tremblay S, Testard C, Inchauspé J, Petrides M (2023) Epiphenomenal neural activity in the primate cortex. bioRxiv 2022.09.12.506984. 10.1101/2022.09.12.506984

Tsutsui-Kimura I, Matsumoto H, Akiti K, et al (2020) Distinct temporal difference error signals in dopamine axons in three regions of the striatum in a decision-making task. Elife 9:1–39. 10.7554/ELIFE.62390

van Elzelingen W, Goedhoop J, Warnaar P, et al (2022) A unidirectional but not uniform striatal landscape of dopamine signaling for motivational stimuli. Proc Natl Acad Sci U S A 119:e2117270119. 10.1073/PNAS.2117270119/SUPPL_FILE/PNAS.2117270119.SAPP.PDF

Van Elzelingen W, Warnaar P, Denys D, et al (2022) Striatal dopamine signals are region specific and temporally stable across action-sequence habit formation. Current Biology 32:. 10.1016/j.cub.2021.12.027

Versprille LJFW, McKenzie C, Amorim FE, et al (2026) Dopamine Dynamics in the Nucleus Accumbens Reflect Confidence in Detecting the Occurrence and Nonoccurrence of Visual Signals in Perceptual Decision-Making. Journal of Neuroscience 46:e1696252026. 10.1523/JNEUROSCI.1696-25.2026

Weiner IB, Freedheim DK, Schinka JA, Velicer WF (2003) Handbook of psychology. Volume 2, Research methods in psychology

Wu T, Hallett M, Chan P (2015) Motor automaticity in Parkinson’s disease. Neurobiol Dis 82:226. 10.1016/J.NBD.2015.06.014

Wu Z, Lin D, Li Y (2022) Pushing the frontiers: tools for monitoring neurotransmitters and neuromodulators. Nat Rev Neurosci 23:257–274. 10.1038/S41583-022-00577-6

Yin HH, Ostlund SB, Balleine BW (2008a) Reward-guided learning beyond dopamine in the nucleus accumbens: The integrative functions of cortico-basal ganglia networks. Eur J Neurosci 28:1437. 10.1111/J.1460-9568.2008.06422.X

Yin HH, Ostlund SB, Balleine BW (2008b) Reward-guided learning beyond dopamine in the nucleus accumbens: The integrative functions of cortico-basal ganglia networks. Eur J Neurosci 28:1437. 10.1111/j.1460-9568.2008.06422.x

Zhou FM, Wilson C, Dani JA (2003) Muscarinic and nicotinic cholinergic mechanisms in the mesostriatal dopamine systems. Neuroscientist 9:23–36. 10.1177/1073858402239588

