## Supplement File for "Differential contributions of striatal dopaminergic pathways to visuomotor conditional learning"

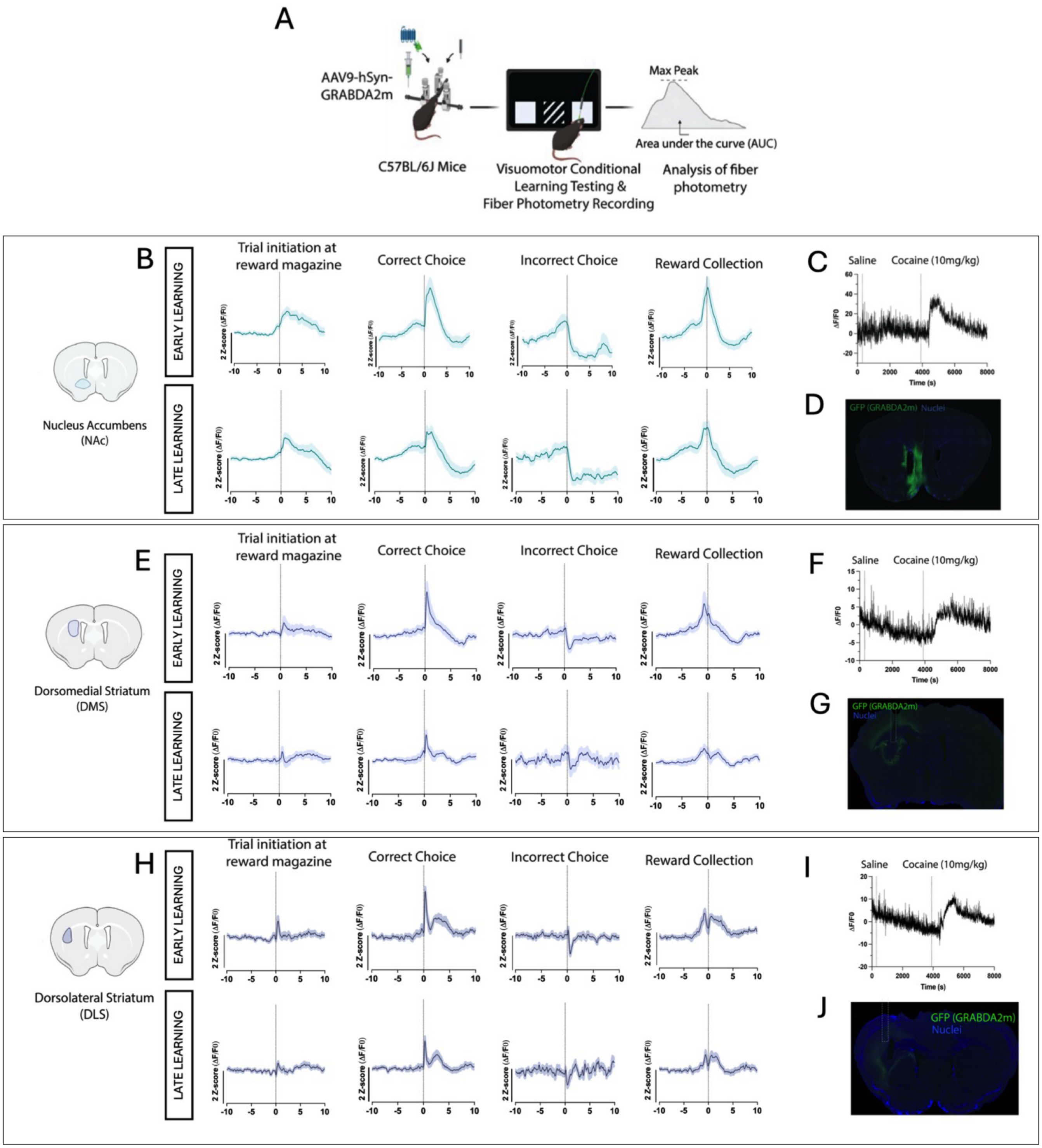


**Supplementary Fig. 1 | Dopamine (DA) signals in striatal regions at early (session 1) and late learning (session 19) aligned to four key task events (Trial Initiation, Correct and Incorrect Choice and Reward Collection).**

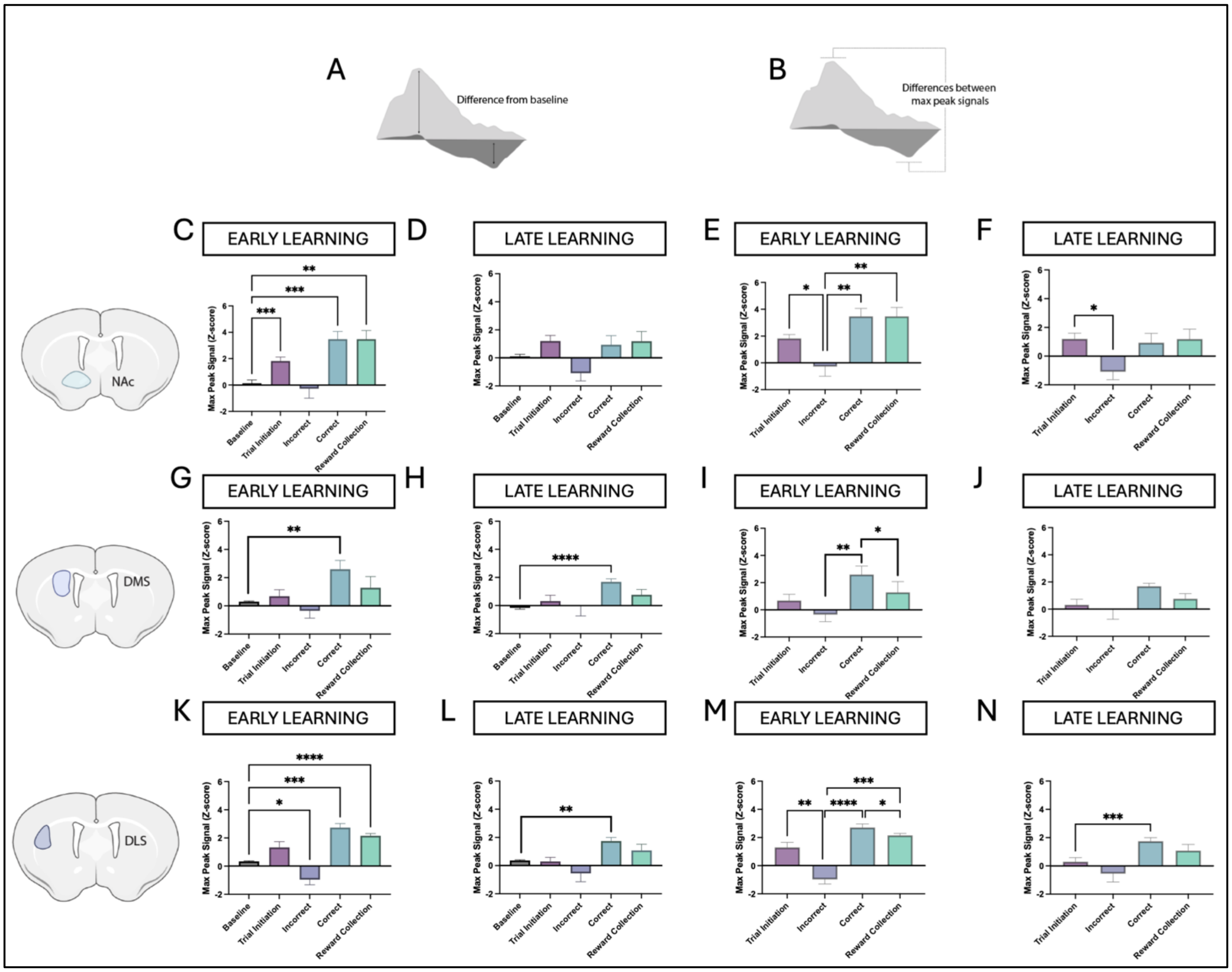


**Supplementary Fig. 2 | *In vivo* dopamine dynamics in the nucleus accumbens, dorsomedial and dorsolateral striatum differ from baseline and from one another most strongly in early relative to late learning.**

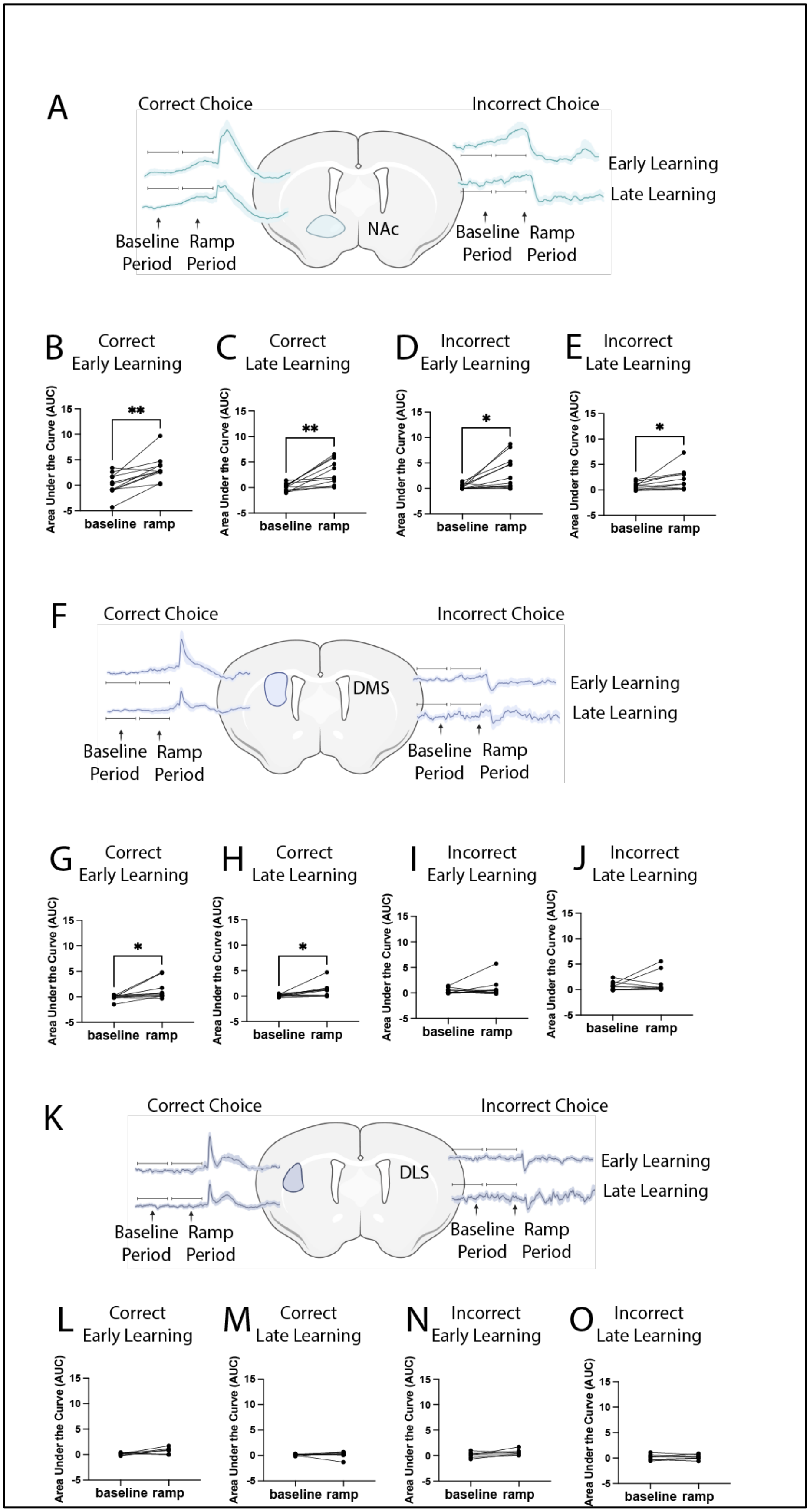
A
Correct Choice
Incorrect Choice
Early Learning
Late Learning
NAc
Baseline Period
Ramp Period
B Correct Early Learning
C Correct Late Learning
D Incorrect Early Learning
E Incorrect Late Learning
Area Under the Curve (AUC)
baseline ramp
**
*
F
Correct Choice
Incorrect Choice
Early Learning
Late Learning
DMS
Baseline Period
Ramp Period
G Correct Early Learning
H Correct Late Learning
I Incorrect Early Learning
J Incorrect Late Learning
K
Correct Choice
Incorrect Choice
Early Learning
Late Learning
DLS
Baseline Period
Ramp Period
L Correct Early Learning
M Correct Late Learning
N Incorrect Early Learning
O Incorrect Late Learning

**Supplementary Fig. 3 | Ramp-like activity preceding choice is most prevalent in the nucleus accumbens, and not present in the dorsolateral striatum.**

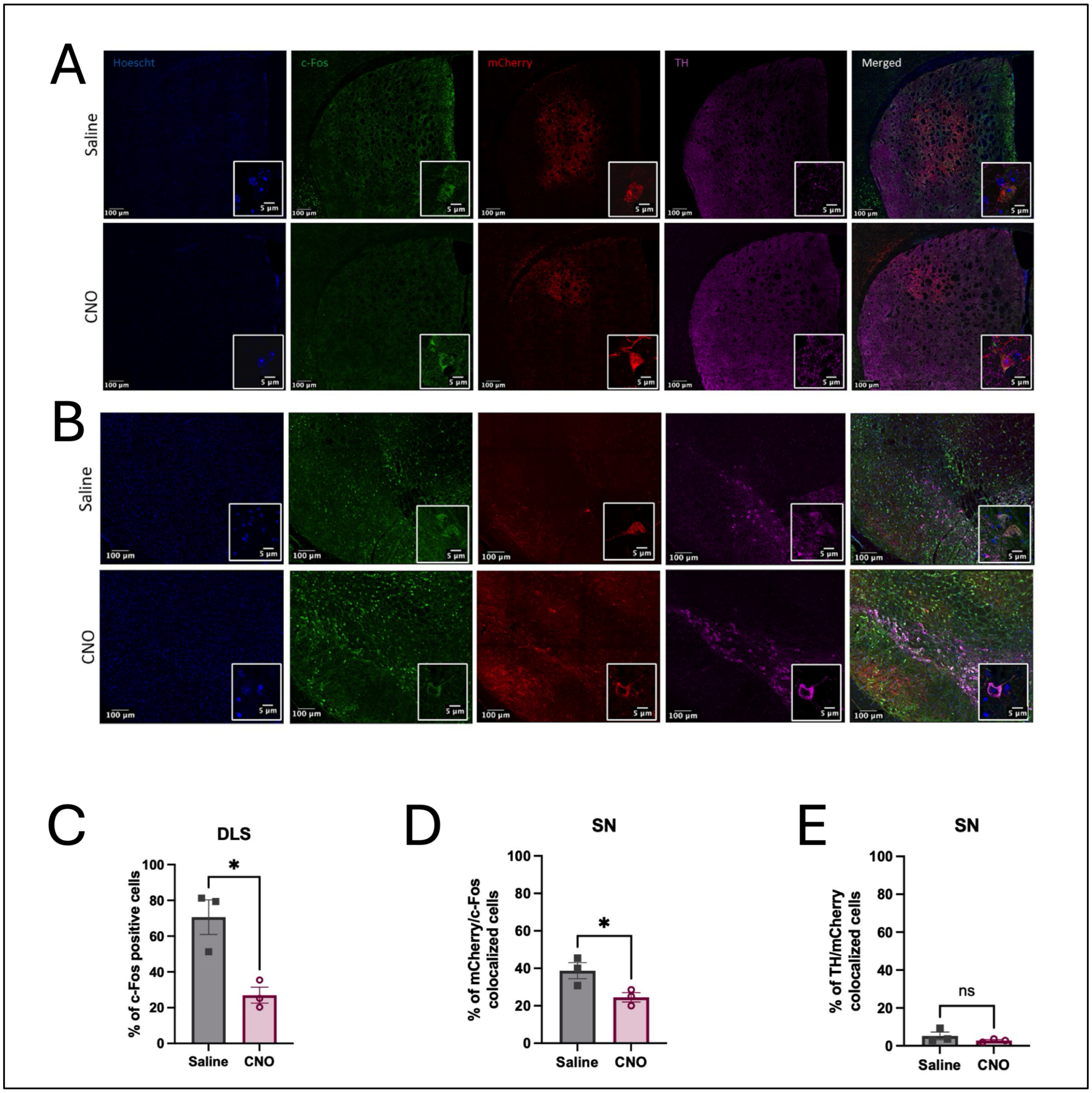


**Supplementary Fig. 4 | Chemogenetic viral infection within the dorsolateral striatum enabled selective expression and inhibition of substantia nigra pars compacta dopamine neurons.**

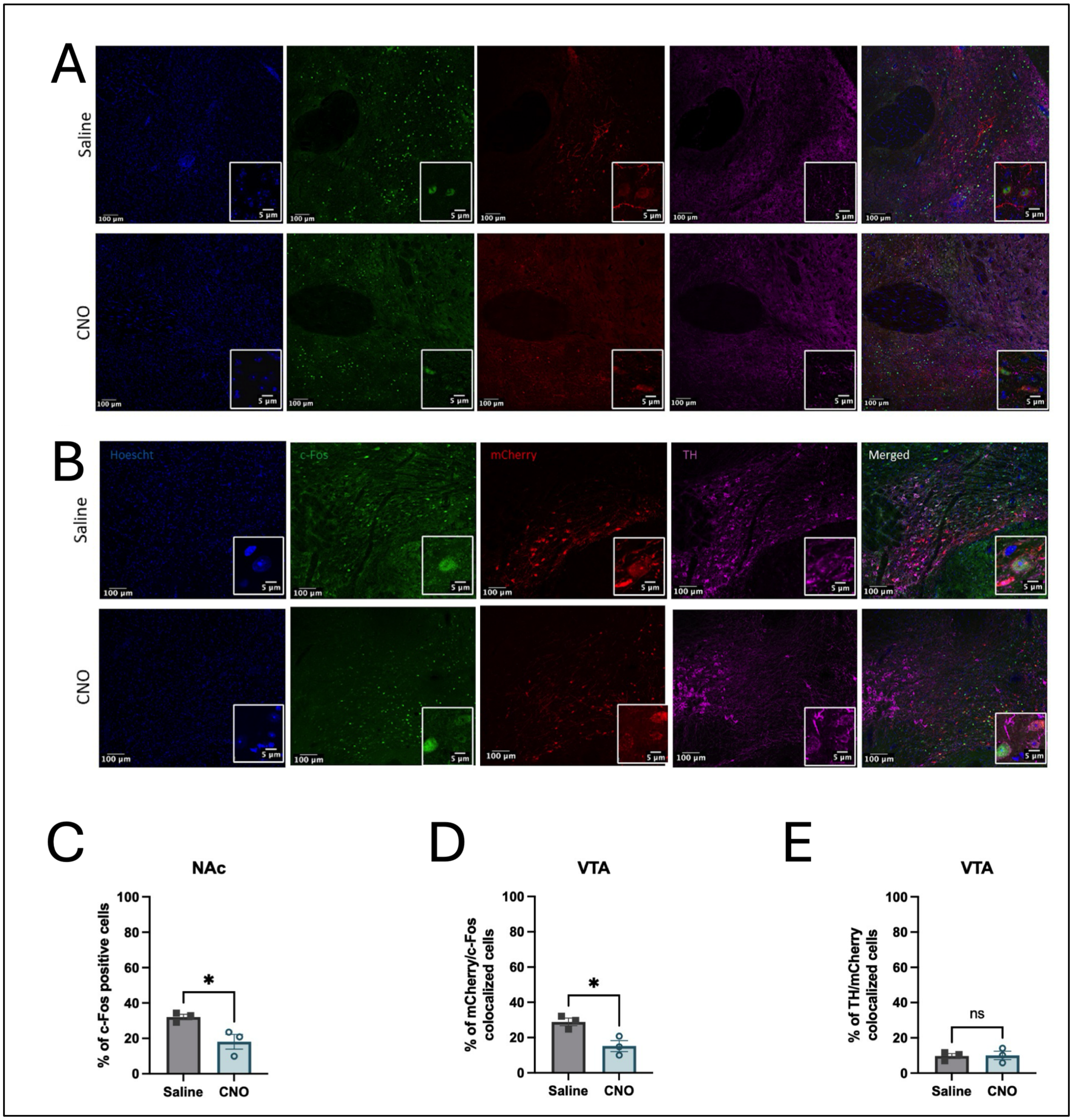


**Supplementary Fig. 5 | Chemogenetic viral infection within the nucleus accumbens enabled selective expression and inhibition of ventral tegmental area dopamine neurons.**

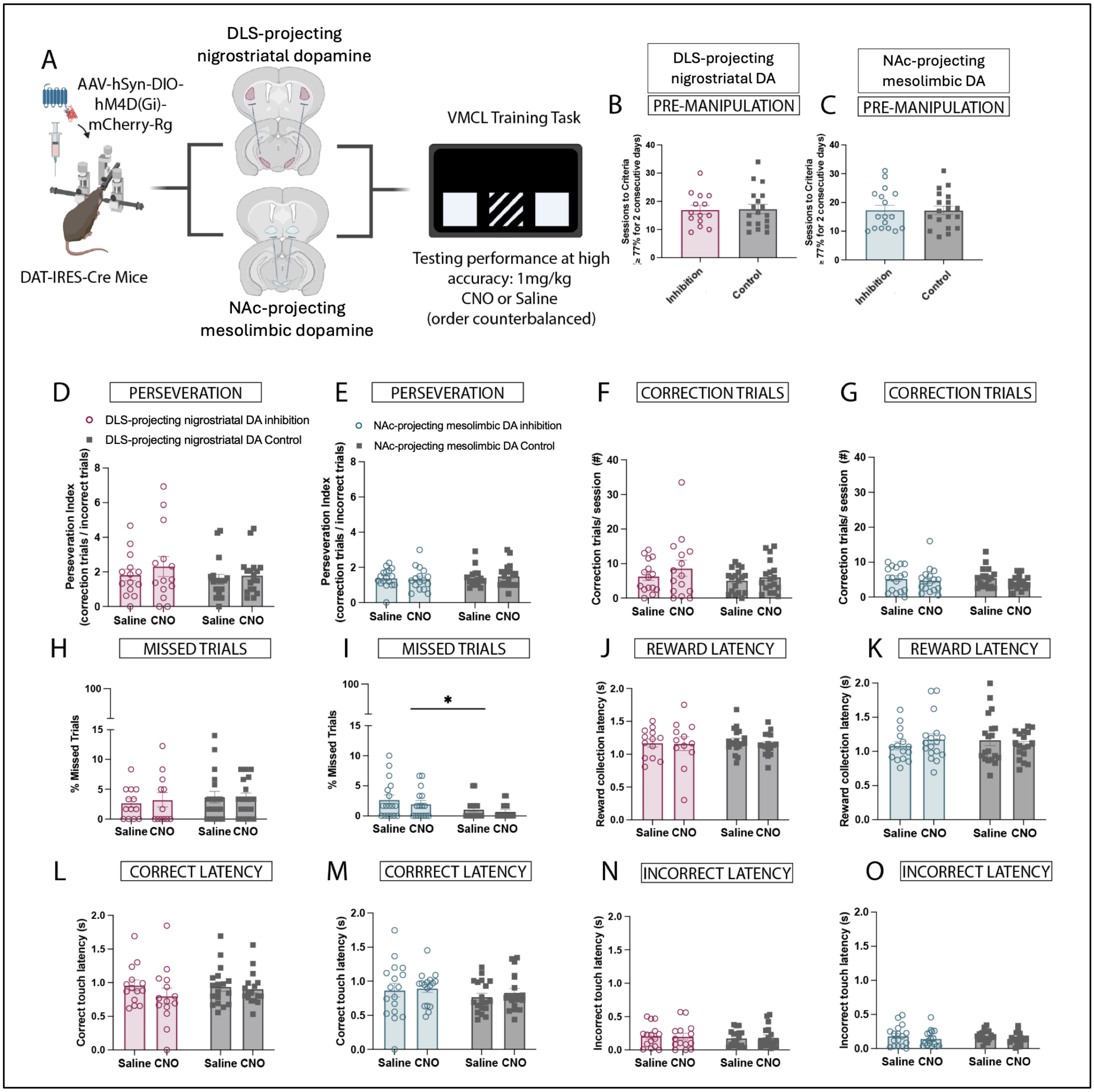


**Supplementary Fig. 6 | Inhibition of DLS-projecting nigrostriatal and NAc-projecting mesolimbic dopamine has minimal impact on expression of previous learning.**

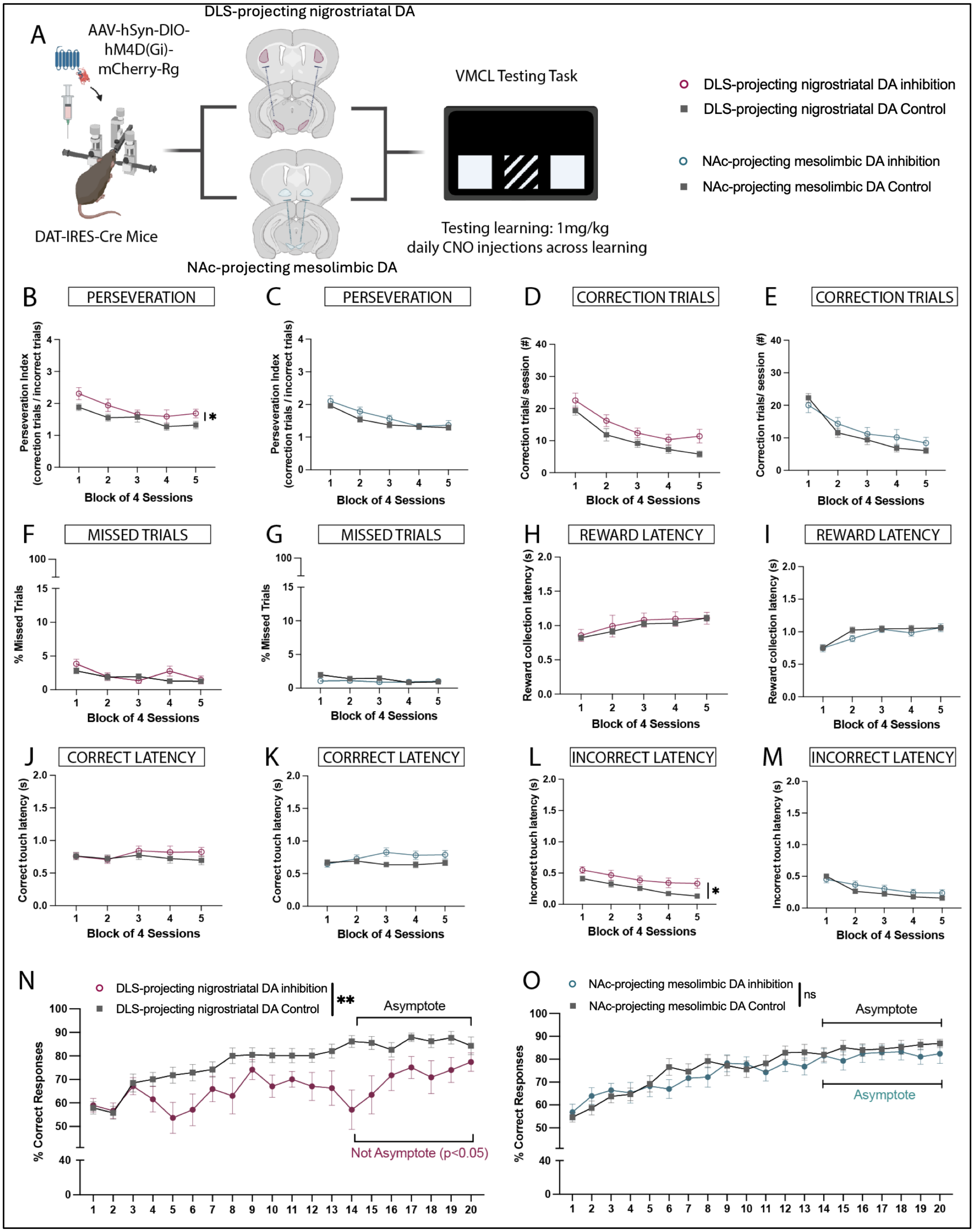


**Supplementary Fig. 7 | Inhibition of DLS-projecting dopamine but not NAc-projecting mesolimbic dopamine led to selective impairment in visuomotor conditional learning.**

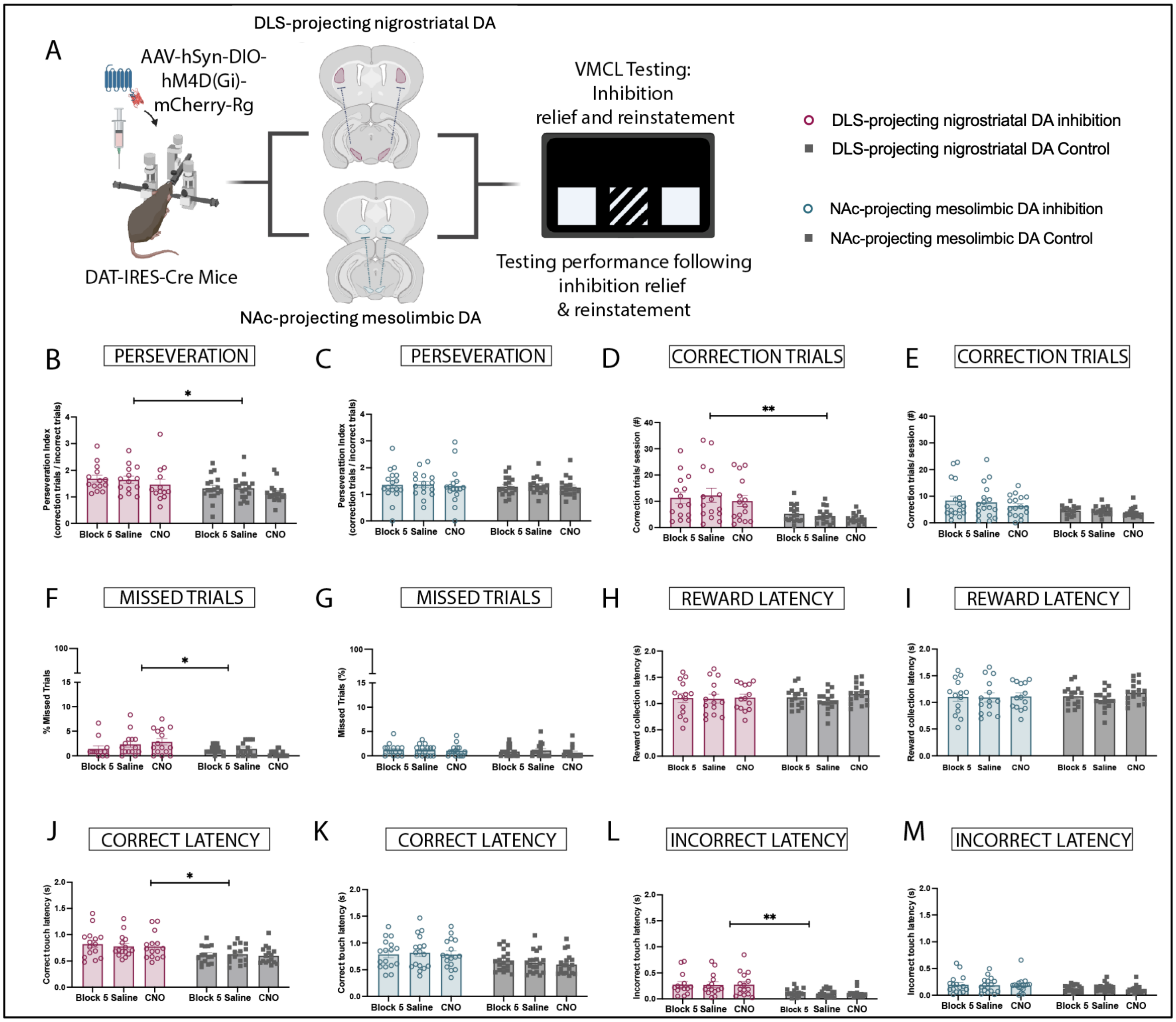


**Supplementary Fig.8 | DLS-projecting nigrostriatal, but not NAc-projecting mesolimbic dopamine inhibition has a long-lasting impact on visuomotor conditional learning test performance.**
